# Brain functional connectivity dynamics in the aftermaths of affective and cognitive events

**DOI:** 10.1101/685396

**Authors:** Julian Gaviria, Gwladys Rey, Thomas Bolton, Jaime Delgado, Dimitri Van de Ville, Patrik Vuilleumier

**Affiliations:** Laboratory for Behavioral Neurology and Imaging of Cognition; Department of Fundamental Neurosciences, University of Geneva, Geneva, Switzerland; Swiss center for Affective Sciences, University of Geneva; Medical Image Processing Lab, Institute of Bioengineering/Center for Neuroprosthetics, École Polytechnique Fédérale de Lausanne (EPFL), Geneva, Switzerland; Institute Department of Radiology and Medical Informatics, University of Geneva, Geneva, Switzerland

**Keywords:** Brain networks, cognitive control, co-activation patterns, emotions, dynamic functional connectivity (dFC), negative affect

## Abstract

Neuroimaging studies have shown carry-over effects on brain activity and connectivity following both emotional and cognitive events, persisting even during subsequent rest. Here, we investigate the functional dynamics of such effects by identifying recurring co-activation patterns (CAPs). Using the precuneus as seed region, we compare carrying-over effects on brain-wide CAPs and their modulation after both affective and cognitive challenges. Female volunteers (n=19) underwent fMRI scanning during emotional induction with sad movie clips, and executive control tasks, each followed by resting periods. Several CAPs, overlapping the default mode, salience, attention, and social cognition networks were impacted by both the preceding events (movie or task) and their valence (neutral or negative), with differential fluctuations over time. Specifically, a modulation of CAPs in posterior cingulate and ventromedial prefrontal cortex was observed after exposure to negatively valenced emotional content and predicted changes in subjective affect. Additionally, CAPs in anterior cingulate cortex and dorsal fronto-parietal areas were induced by cognitive control in a negative, but not neutral context, and amplified by the task difficulty. These findings provide new insights on the anatomical organization and temporal inertia of intrinsic functional brain networks, engaged by transient emotions and presumably involved in subsequent adaptive homeostatic processes.

## INTRODUCTION

Emotions are an essential component of mental life and adaptive behaviors, with profound impact on our perception, memories, actions, and thoughts. In particular, negative affect may impair the efficiency of high-level cognitive processes and reduce cognitive control (Hilt et al. 2014). Further, experiencing negative emotions has been associated with cognitive inflexibility (Davis and Nolen-Hoeksema 2000) and difficulties in mental switching (Piguet et al. 2010, 2016). A better understanding of such influences of affective states on cognitive control and mental functions is therefore warranted to elucidate mechanisms by which emotions and moods can steer behavior in adaptive or maladaptive manners, in both health and disease.

Recent neuroimaging research has begun to unveil the neurobiological bases of how affective signals can induce changes in cognition and brain functioning. Different approaches have been employed to characterize these effects (Okon-Singer et al. 2015). On one hand, standard (univariate) brain imaging approaches allow pinpointing the role of particular brain regions in responding to specific categories of emotions (e.g., fear) and triggering concomitant changes in the activity of other brain areas implicated in cognition, memory, or action (Vuilleumier et al. 2004; Okon-Singer et al. 2007; Vuilleumier and Huang 2009; Etkin et al. 2011). For example, increased amygdala activity can directly regulate the recruitment of sensory (e.g., visual) cortices, hippocampus, or motor areas (Dolan and Vuilleumier 2010; Pourtois et al. 2010). On the other hand, functional brain changes evoked by emotions are widely distributed and can be defined in relation to broader networks (Pourtois et al. 2013), including those associated with domain-general core functions such as the frontoparietal attention control (FPCN), salience (SN), or default mode networks (DMN) (Göttlich et al. 2017). Importantly, in both approaches, emotion and cognition are recognized as deeply intertwined and subserved by overlapping neural systems that jointly contribute to adaptive (or maladaptive) behaviors (Scherer 2009; Lindquist and Barrett 2012; Braver et al. 2014; Okon-Singer et al. 2015; Touroutoglou et al. 2015; Pessoa and McMenamin 2017).

Recent studies also showed that emotions can lead to lingering changes in brain activity that persist long after the eliciting episodes, even during resting state, including changes in the functional connectivity of DMN areas (such as precuneus or post-cingulate cortex) with amygdala after fearful or joyful events (Eryilmaz et al. 2011), or with striatum and orbitofrontal cortex after reward or loss events (Eryilmaz et al. 2014). In turn, these “recovery” related changes may affect subsequent affective processing in face recognition tasks (Qiao-Tasserit et al. 2017) or social cognition (Qiao-Tasserit et al. 2018). These lingering changes may also be associated with maladaptive emotional inertia and ruminative processing observed in several psychopathological conditions (Kuppens et al. 2010). However, it remains unresolved whether post-emotion changes in resting brain activity are specific to affective episodes or whether they may also occur after other behavioral challenges (e.g. difficult task demands).

In addition, previous imaging approaches to emotion-induced changes (Eryilmaz et al. 2011) remain limited in that they provide a relatively static view of brain activity patterns. Most functional connectivity measures between regions of interest and network-based analyses at the whole-brain level derive from a correlation of activation time-courses in different areas averaged over long time periods (often several minutes). This might be insufficient for fully characterizing the temporal neural dynamics by which emotional and cognitive systems interact in the brain. Growing evidence suggests that functional connectivity (FC) dynamically varies over time, not only as a function of different task and motivational variables but also during spontaneous resting sate (Betzel et al. 2016, 2017, Cole et al. 2016a, 2016b). For instance, a given brain region might have more frequent but transient moments of connectivity, with a few other regions during a particular emotion episode, or during a recovery period following this emotion. But this might not be captured by an average correlation of their time-courses during the fMRI scanning time. Such limitations can partly be circumvented by new methods designed to track the temporal dynamics of functional connectivity (dFC) (Chang and Glover 2010; Hutchison et al. 2013; Preti et al. 2017), allowing for a finer-grained dissection of brain activity configurations as it evolves at different time scales (Karahanoğlu and Van De Ville 2017).

Among dFC approaches, a recent method has been proposed to regard fMRI volumes at individual time points, instead of time courses, as the basic units of analysis (Tagliazucchi et al. 2012), and to extract recurring co-activation patterns (CAPs) among whole-brain voxels (Chang et al. 2016) in order to determine their fluctuations over time (Liu, Zhang, et al. 2018). The CAPs characterize instantaneous configurations of activity across distributed networks, without the need of parcellation or atlasing into pre-defined regions. This approach is particularly valuable for studying the temporally dynamic nature of brain connectivity associated with emotion-cognition interactions because it allows for whole brain analysis without relying on a priori, fixed networks and provide measures of transient fluctuations in the configurations of networks that may anatomically overlap in parts. In addition, the CAP approach has a low sensitivity to noise, thanks to an exclusion of time-points with small activity amplitude, which are more likely to contain noise rather than reflect information processing. Accordingly, applying the CAP methodology may significantly improve network analysis as compared with more classical strategies (Li et al., 2014b).

In the current study, we applied the CAP approach to fMRI and collected concomitant behavioral and psychophysiological data during a novel paradigm, in which we probed for an impact of both (negative) emotion and (difficult) cognitive challenges on subsequent spontaneous brain activity at rest. Cognitive control is defined as the flexible regulation of behavior in the pursuit of internal goals (Egner 2007; Helfrich and Knight 2016), and commonly assessed by Stroop and Flanker tasks where responding to goal-relevant targets is tested in the presence of distracting information (Botvinick et al. 2001; Schmidt et al. 2015). Negative emotions have been shown to alter cognitive control in such tasks at both the behavioral (van Steenbergen et al. 2010; Schuch et al. 2012; Brown et al. 2014; Schuch and Koch 2014) and neural levels (Egner et al. 2008; Etkin et al. 2011; Braem et al. 2013; Brown et al. 2014; Song et al. 2017; see also van Steenbergen 2015 for a review). Here we built on previous work showing an impact of negative affective episodes on subsequent brain resting state (Eryilmaz et al. 2011) and cognitive control performance (Schuch and Koch 2014; van Steenbergen et al. 2015), by probing for modulations of intrinsic functional brain connectivity and dynamics at the whole brain level following both affective and cognitive challenges. By combining both stimulation types and leveraging the CAP methodology, we aimed to uncover, first, how the recruitment of cognitive control processes may be influenced by negative affect, and second, how resting state connectivity patterns are modified by negative affect alone or by associated cognitive effort. In addition, while previous work focused on highly arousing emotions such as fear (Eryilmaz et al. 2011), here we examined the effect of sadness which is more relevant to mechanisms of mood disorders such as depression (Mayberg et al. 2005). Specifically, transient emotion episodes were induced by viewing brief movies with sad content, after which participants either rested immediately or performed a difficult cognitive control task (Stroop or Flanker) before resting. Brain activity was measured with fMRI in all phases (see Figure. 1A). Based on previous behavioral (van Steenbergen et al. 2010; Schuch and Koch 2014; Gendolla et al. 2015; van Steenbergen 2015) and neuroimaging work (Eryilmaz et al. 2011; van Steenbergen et al. 2015; Borchardt et al. 2017; Song et al. 2017; Young et al. 2017), we predicted that negative affective contexts would cause distinctive changes among functionally integrated brain-wide networks at rest, including the DMN, SN, FPCN, and possibly other limbic networks (e.g., centered on amygdala and insula), with particular temporal fluctuations and distinctive distribution as a function of the preceding events. This allowed us to investigate several key questions including: (1) whether negative affect episodes produce lingering changes in subsequent brain activity and connectivity at rest, (2) whether these neural changes are also sensitive to the cognitive demands of executive control tasks, and (3) whether performance in executive control tasks themselves is altered by negative affect. Altogether, our results provide new insights about the neural architecture and temporal dynamics of brain circuits mediating emotion-cognition interactions.

## MATERIALS AND METHODS

### Participants

Thirty-four right-handed, French-speaking female volunteers (mean age= 23.2 ± 4.3) were contacted via posters and online advertisement. They all provided written informed consent according to the Geneva University Hospital Ethics Committee. Only female participants were recruited because pilot testing suggested stronger emotional induction in women, compared to male, particularly with the movie clips used here (see Supplementary methods). Controlled inclusion criteria were: no history of neurological and psychiatric diseases; known menstrual phase and no contraceptive method to rule out hormonal effects on emotional processing (Protopopescu et al. 2005) and functional connectivity (Petersen et al., 2014)(Pletzer et al., 2016); no major head mouvement during the scanning sessions (<1mm in all axes). The final sample consisted of nineteen subjects. Scanning was scheduled for each participant in the early stage of the menstrual follicular phase, when the levels of estradiol and progesterone hormones are moderate (Endicott 1993; Andreano et al. 2018). Nicotine and caffeine consumption were prohibited 10 hours before scanning.

### Experimental design

Upon arrival, the participants filled in the French version of two mood questionnaires (PANAS and STAI). Next, they received general instructions about the experimental session and trained on the two cognitive tasks (25 trials from the Flanker and 25 from the Stroop). This was followed by fMRI scanning for approximately 62 minutes (see Fig. 1 for schematic overview). It comprised a first 5 min resting state (baseline), followed by an emotional context period with movies containing either neutral or negative emotional scenes (order counterbalanced across participants). Each emotional context period included two video clips with content of the same valence (5 min per clip), the first followed by another resting block with open eyes (5 min), the second followed by a cognitive control task (6 min) and then a final resting block (5 min). The cognitive control task consisted of either a Stroop or Flanker paradigm (counterbalanced across emotion contexts and participants). Finally, post-experiment mood questionnaires were filled again, in addition to a brief questionnaire about the movies. Detailed information on movies, affective ratings (PANAS and STAI), and recording of physiological measures (pupillometry) is provided in the Supplementary methods.

**Figure 1.**
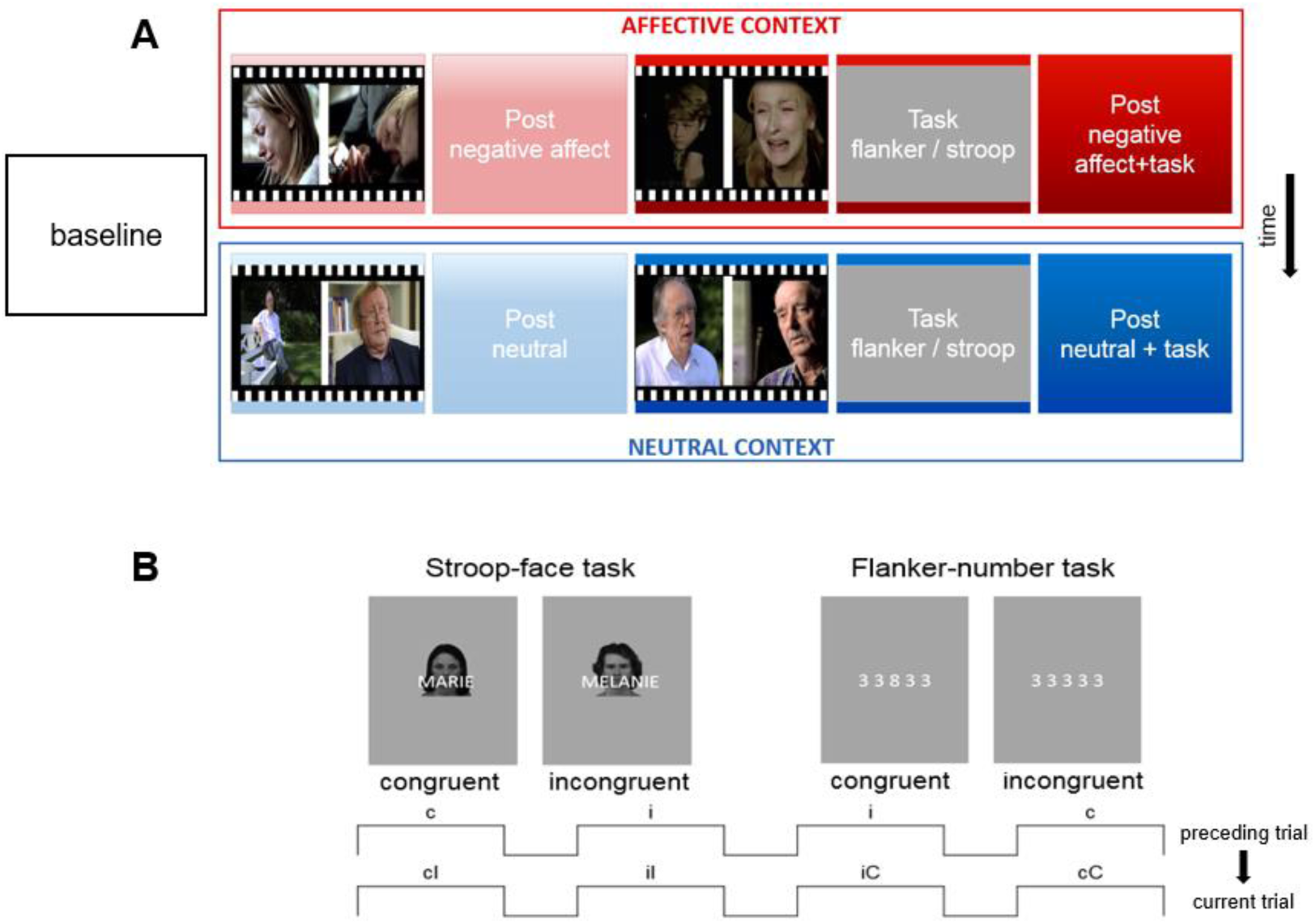
**A)** Paradigm design illustrating the sequence of experimental events. Sad or neutral movies were followed by a rest period, after which a second movie block with the same valence was followed by a cognitive task and then another rest period. A baseline rest period was presented prior to movies in all conditions. The order of the affective context was counterbalanced across participants. The Stroop and Flanker tasks were counterbalanced across participants and affective contexts. **B) Top**: Illustration of stimuli in the Stroop task (left) and Flanker task (right). **Bottom:** Illustration of trial sequence in each cognitive task. It combined congruent (C) and incongruent (I) trials that could be preceded by either the same or opposite condition (C or I trials), in a semi-random but balanced order. This resulted in 4 trial types, allowing subsequent behavioral analysis according to both current compatibility (indicated by upper-case letters C and I) and compatibility of the preceding trial (indicated by lower-case letters c and i).

### Cognitive control tasks

To probe for the functional aftermath of cognitive control demands on subsequent resting state, and its modulation by affective context, following the work of Schuch and Koch (2014), we used two cognitive tasks known to engage selective attention, generate reliable interference effects with frequent errors, and are sensitive to affective conditions (see fig. 1B). One task was a Flanker-number paradigm with digits as stimuli (Eriksen and Eriksen 1974; Hübner et al. 2010; Schuch and Koch 2014); and the second a Stroop-like paradigm with superimposed face–name stimuli (Stroop 1935; Egner and Hirsch 2005; Schuch and Koch 2014). Multiple factors led us to choose these two tasks. First, previous studies reported high sensibility of both tasks to negative emotional priming (Mayr et al. 2003; Hommel et al. 2004; Egner 2007; Mayr and Awh 2009; Schuch and Koch 2014). Second, both tasks produce reliable response conflicts, with the possibility to present many trials, with minimal learning, and both allow measuring post-conflict adaptation effects on RTs across successive trials (Macleod 1991; Verguts et al. 2011). Third, our pilot results yielded very similar interference effects in both tasks, with and without emotional modulation (see results in Results section), allowing us to balance them across conditions and participants to avoid systematic order effects. In turn, observing similar behavioral results and emotional influences on performance across two paradigms would provide evidence for the generality and robustness of our finding (Schuch and Koch 2014). It is also important to note that our index for assessing any emotional influence on cognitive control performance was the interference magnitude (RT on congruent vs. incongruent trials) within a given task; hence trials from one task (e.g. Flanker) were not compared to those from the other task (e.g. Stroop). Detailed information about these tasks is provided in the Supplementary methods.

### MRI data acquisition

MRI data were acquired with a 3T scanner (Siemens TIM Trio) at the Brain & Behavior Laboratory, University of Geneva. A multiband-sequence was implemented, with a voxel size of 2mm isometric and a TR of 1.3s (see in supplementary material for details on data acquisition and preprocessing).

### Co-activation pattern (CAP) analysis

#### Dynamic functional connectivity of DMN nodes defined by CAPs

In a nutshell, CAP analysis allows decomposing the BOLD signal at rest into multiple spatial patterns that express the ongoing dynamic organization and reorganization of brain activity over time (see Liu et al, 2018 for comprehensive overview; and Preti et al., 2017 for application within the framework of dynamic functional connectivity). A key feature of this method is the selection of a seed region of interest (ROI), with which the rest of the brain co-activates under different spatial configurations at specific time points. Here we selected the bilateral precuneus as seed, based on its central contribution to the default mode network (DMN; Utevsky et al., 2014) as well as its role in self-consciousness (Vogt and Laureys 2005), memory (Cavanna and Trimble 2006), and attentional flexibility (Piguet et al. 2016). Moreover, the precuneus was also found to show higher activity during affective vs. neutral movies in our main GLM analysis (see below and Supplementary Figure S1.A). This seed region comprised 486 voxels, covering the mid dorsal, central, and ventral central sectors of the precuneus. The time-course of this ROI was then z-scored and thresholded to define time points at which fMRI frames were selected for subsequent analysis. These frames were submitted to a first temporal clustering step to define the number of clusters (CAPs) to retain for final classification. For this purpose, we implemented a consensus clustering method (Monti et al., 2003), which provides quantitative evidence for evaluating the number and membership of possible clusters within a dataset. Apart from the seed ROI and the number of clusters, input data for the CAP analysis also included a gray-matter segmentation mask from each participant’s brain (see Supplementary methods, preprocessing section), the dimension and spatial position of data, individual framewise displacement information, and the BOLD fMRI data itself. The BOLD time series were extracted and Z-scored voxelwise across the whole gray-matter brain volume. Data from all resting conditions were pooled together in order to compute CAPs in one single analysis and statistically compare the different resting conditions, so as to identify changes in the main spatiotemporal features of these CAPs as a function of the preceding affective and task events.

#### Statistical analysis of CAPs across conditions

Data were analyzed using the “R” statistical software version 3.6.0 (RStudio Team, 2018). We computed mixed linear models (lmms) with the “lmerTest” (Kuznetsova et al., 2017) and “lme4” (Bates et al. 2014) packages, focusing on the spatiotemporal features of CAPs observed during the resting periods. P-values for fixed effects were obtained by Satterthwaite’s method, and sums of squares for the analyses of variance (ANOVAs) were computed with the type II method (Herr 1986). Due to their flexible framework, lmms are more efficient to model repeated measures with missing data (Garrett et al. 2011). A preliminary inspection on the plot residuals did not reveal any obvious deviation from homoscedasticity or normality (Suppl. Figure S5).

Our statistical analysis of CAPs comprised three stages. A first set of lmms (one for each CAP) examined their occurrence variance across our 5 main experimental rest conditions (baseline, post affective, post task affective, post neutral, post task neutral). To account for non-independence of repeated data, individual participants and “occurrence rates” of each CAP were modeled as random factors, while the rest conditions were introduced as fixed factors. Only those CAPs showing a significant main effect in this stage were considered for further analysis (Figure 2).

**Figure 2.**
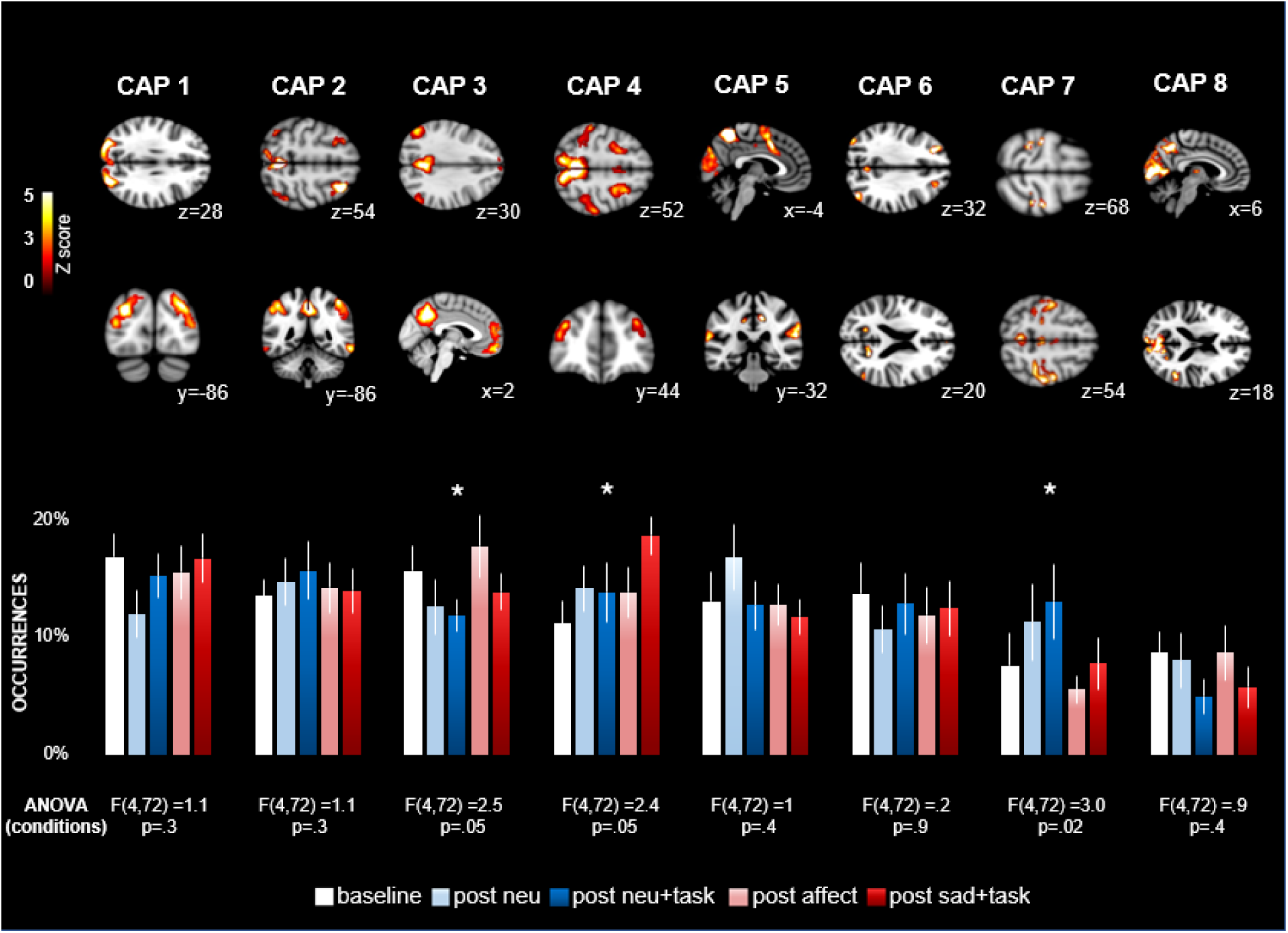
Illustration of 8 CAPs found to be most reliably active during rest periods. Up: Spatial maps of CAPs denoting brain areas showing positive co-activation with the bilateral precuneus seed, across all rest conditions. Bottom: Occurrence rates for each CAP denoting their temporal fluctuation in the different resting conditions. A LMM analysis of occurrence rates for each CAP revealed that 3 out of the 8 clusters presented significant differences between conditions (Coan and Allen 2007).

Next, a second set of lmms was computed on the temporal metrics of selected CAPs (entry rates and duration). This second stage identified significant differences in each CAPs, across all conditions, for all three temporal metrics. Finally, the third stage of analysis consisted of a new set of lmms in which the baseline rest condition was excluded, now including only the 4 critical experimental contexts defined by emotion (neg. affect vs neutral) and preceding event (post film vs. post task) as fixed factors. This third stage of analysis allowed for evaluating main effects of preceding task conditions (movie vs cognitive control) and emotional context (sad vs neutral) separately, as well as their interactions, for each temporal metric of each of the CAPs selected from the second stage.

We also explored possible correlations of the different temporal metrics of CAPs during resting conditions with measures of subjective emotional state (PANAS) and behavioral indices of interference in the cognitive control tasks. For this purpose, we computed Pearson’s correlation coefficients, followed by a multivariate permutation test, a nonparametric approach to control for multiple significance testing (Yoder et al. 2004). A total of 10.000 permutations were implemented, in order to obtain an accurate approximation of the of the exact probability (p) value, followed by correction for multiple comparisons. Finally, paired T-test contrasts in SPM12 were performed to compare spatial CAPs between conditions (i.e., “post affective task vs. post neutral task”), and their relation to affective and behavioral scores.

#### CAPs time series analysis across resting periods

As CAPs allow for capturing dynamic fluctuations in brain activity, we evaluated temporal changes in the occurrence rates of CAPs of interest (No 3, 4, and 7) over the five minutes of each resting period. To do so, we segmented the temporal transition probability matrix (i.e., the matrix describing the probability of switching from one CAP to another during the five minutes of each resting condition) in successive, equal time-bins of one minute. We then performed a chi-square analysis to assess the frequency distribution of occurrence rates for each CAP across each of the resting conditions, in both the neutral and affective contexts (five time-bins per condition).

## RESULTS

### Physiological, behavioral, and neural indices of effective emotional induction by movies

Movies with negative valence induced distinctive patterns in both behavior and brain measures, compared to neutral movies. Subjective affective ratings of the movie clips confirmed a reliable change in emotional experience with more negative valence and higher arousal after “negative affect” compared to “neutral” movies (Suppl. results and Suppl. Figure S1.C). Pupillary diameter was also significantly larger in the affective context (Suppl. Figure S1.B). In addition, both the STAI and PANAS scores differed between the pre and post scanning measures. These data validate an effective manipulation of emotional state by our movies with a negative valence (see also supplementary results and Suppl. Figure S1.D).

Differential brain activity during the negative vs. neutral movies was also demonstrated by direct contrasts between conditions using a GLM-based flexible factorial analysis in SPM (see Suppl. Methods and Suppl. Table S1). As expected, this analysis yielded higher activations (FWE p<.05) in bilateral visual areas, temporo-parietal junction, and precuneus during negative affect movies (see Suppl. Figure S1. A. Suppl. Table S1), relative to neutral, consistent with greater engagement of perceptual processes, social cognition, and access to self-relevant or autobiographical memory information in response to sad scenarios in these movies (Elman et al. 2013; Chen et al. 2017; Göttlich et al. 2017; Pan et al. 2018). Conversely, neutral movies produced greater activation in parietal and frontal cortices, associated with voluntary attention, working memory, and more effortful cognitive processing (Crottaz-Herbette et al. 2004; Mohr 2006; Mayer et al. 2007) (see results in Supplementary Table S1).

### Affective modulation of cognitive control tasks

We first verified that an attentional interference effect was reliably elicited in both cognitive tasks (Stroop and Flanker), by comparing reaction times and error rates on incongruent vs congruent trial types in each task separately (I>C). Data are shown in Suppl. Figure S2.A). We computed the magnitude of both the main interference / congruency effect (CE) and the congruency sequence effect (CSE; Egner 2007) across the two emotion context conditions, which was then submitted to a lmm-based ANOVA with separate factors for affective context (post neutral vs post neg. affect), current trial type (xC and xI), and previous trial type (Cx, Ix). This analysis for RTs yielded a significant interaction between current trial type and emotion context (F (1,126) = 4.9, p < 0.05); and between current and previous trial types (F (1,126) = 46.1, p < 0.001); but no triple interaction (F (1,18) = .002, p <1) (see Table S2). The same ANOVA on error rates yielded only a significant effect of current trial (interference) (F (1,126) = 14.5, p < 0.001), across both tasks, and regardless of affect context (Table S2). Importantly there were no main effects or interactions involving the task, confirming that both our Stroop and Flanker paradigms elicited generally similar cognitive control performances.

**Table 2.** Results of LMMs for the analyses of occurrences (models O2 CAP3; O2 CAP7), entry rates (models E2 CAP3; E2 CAP7), and durations (models D2 CAP4), comparing main effects of the two emotional contexts (neg. affect vs neutral) and the two preceding events (movie alone vs movie + cognitive task). Abbreviations: B= Baseline; N= post neutral; NT=post neural + task; A= post negative affect; AT= post negative affect + task. ^.^ p>0.1; * p>0.05; ** p>0.01; ***p>0.001.

| Model | ANOVA | | | Estimates $\beta$ (SE) | | | | LS means difference |
| --- | --- | --- | --- | --- | --- | --- | --- | --- |
|  | Context | Event | Context x Event | N | NT | A | AT |  |
| O2 CAP3 | F(1,54)=5.8. p<.01 | (1,54)=2.4. p<.1 | (1,54)=1.3. p<.1 | 12.5 (2.0) | 11.8 (2.0) | 18.5 (2.0) | 14.0 (2.0) | A > N *<br>A > NT ** |
| O2 CAP7 | F(1,54)=10.6. p<.01 | (1,54)=1.3. p<.1 | (1,54)=1.3. p<.1 | 11.2 (2.6) | 13.0 (2.6) | 5.5 (2.6) | 7.7 (2.6) | NT > A **<br>N > A * |
| E2 CAP3 | F(1,54)=8.0. p<.01 | F(1,54)=0.8. p<.1 | F(1,54)=2.9. p<.1 | 11.3 (1.3) | 13.8 (1.3) | 17.3 (1.3) | 13.8 (1.3) | A > N **<br>A > NT * |
| E2 CAP7 | F(1,54)=13.5. p<.001 | F(1,54)=2.4 p<.1 | F(1,54)=1.1. p<.1 | 12.1 (2.2) | 12.9 (2.2) | 5.6 (2.2) | 9.2 (2.2) | N > A **<br>TN > A ** |
| D2 CAP4 | F(1,54)=7.6. p<.01 | F(1,54)=0.04. p<.1 | F(1,54)=2.5. p<.1 | 3.1 (.3) | 2.6 (.3) | 3.5 (.3) | 4.1 (.3) | AT > N *<br>AT > NT **<br>A > NT * |

These results indicate a robust modulation of interference effects / cognitive conflict (I>C) by negative affective state (van Steenbergen 2015; Song et al. 2017). To further clarify these effects, we calculated a differential CE index (RT to current incongruent minus current congruent trials), which showed a significant increase in the negative affect context compared to the neutral one (F= (1,126)=19.9 p<.05; Figure S1D) but no difference between the Stroop and the Flanker tasks (F (1,18) = 1.3, p > .1; see Figure S2.B). These data confirm that cognitive conflict during attentional control (incongruence between target and distracter information) is amplified by a negative affective context. In contrast, cognitive adjustment (change between current and previous trials) was not modified by affective context (F (1,126)=.002 p>1). While the interaction between current and previous trial types replicated prior work (Botvinick et al. 2001; Egner 2007).

#### Affective modulation of brain co-activation patterns (CAPs) at rest

We next turned to the main goal in the current study concerning the impact of both negative emotions and cognitive control demands on subsequent brain activity and dynamic connectivity at rest. To this aim, we applied the CAP methodology to determine how resting state networks were differentially modulated following exposure to movies alone (post negative affect and post neutral) and exposure to both movies and cognitive control tasks (post negative affect+task and post neutral+task). Clustering results from the CAP analysis were submitted to a consensus selection procedure (Supplementary Figure S3) that yielded K=8 as the optimal number of distinct brain maps in terms of replicability across all resting blocks.

These 8 clusters constitute the most dominant configuration of co-activations over the whole brain during rest in our data set, and are illustrated in Figure 2. Overall, these 8 CAPs exhibited highly plausible functional organization and remarkable similarity with conventional resting-state networks (Seeley et al. 2007; Andrews-Hanna et al. 2010; Dixon et al. 2018).

To identify those CAPs whose presence differed between emotional contexts, we performed a first set of lmms (see Methods section) comparing the occurrence rates of each CAP across conditions. Results showed significance differences for three networks: CAP3, CAP4, and CAP7. These three co-activation clusters were then used for further analysis.

CAP3 exhibited co-active patterns in areas overlapping with the typical default mode network (DMN) including posterior and anterior midline regions, together with co-deactive patterns in areas associated with the salience network (SN) including anterior cingulate and insular regions (Figure 3A, Table 3).

**Figure 3.**
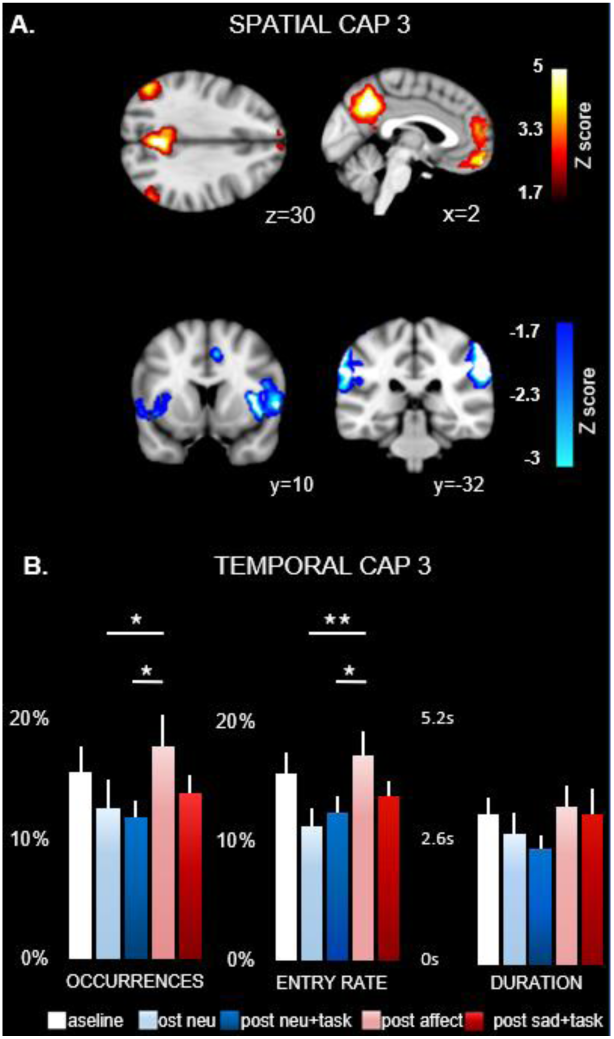
**A)** Temporal analysis of CAP3: the occurrence rate (left), entry rate (middle), and duration (right), averaged across all the participants, is shown for each experimental condition. Stars indicate significant differences among conditions, in a mixed model-based ANOVA analysis (see Results section and table 3). Error bars indicate standard errors of the means (SEM). **B)** Spatial analysis of CAP3. Hot colors (above) represent areas with transient positive (co-)activation with the precuneus seed, while cold colors (below) represent areas with transient negative (de-) activation with precuneus ((see Supplementary table S7 for a further description)). Slice coordinates are provided in MNI space. Brain regions were observed at Z =< −1.7 (p<.05).

**Table 3.** Activity peaks in the spatial cluster maps obtained for CAP3, CAP4, and CAP7. Regions with positive Z-score co-activated with the precuneus seed at Z>1.7 (p<.05). Conversely, brain regions with negative Z-scores were co-deactive in relation to the time-course of precuneus at Z =< −1.7 (p<.05).

| Region (MNI) | x | y | z | K> | Z max. |
| --- | --- | --- | --- | --- | --- |
| **CAP 3** |  |  |  |  |  |
| Angular gyrus | -54 | -68 | 16 | 1226 | 4.1 |
| Angular gyrus | 56 | -62 | 28 | 899 | 3.5 |
| Medial frontal gyrus | -2 | 54 | -16 | 715 | 3.5 |
| Insula | -40 | 2 | 2 | 297 | -2.2 |
| Insula | 36 | 18 | 4 | 736 | -2.5 |
| Supramarginal gyrus | -64 | -26 | 30 | 552 | -2.8 |
| Supramarginal gyrus | 62 | -28 | 40 | 736 | -2.8 |
| Middle cingulate gyrus | 8 | 12 | 38 | 58 | -1.7 |
| **CAP 4** |  |  |  |  |  |
| Superior frontal gyrus | -22 | -2 | 64 | 699 | 2.8 |
| Superior frontal gyrus | 28 | 6 | 62 | 716 | 3.2 |
| Precuneus | 12 | -70 | 54 | 435 | 3.3 |
| Precuneus | -8 | -68 | 56 | 693 | 2.9 |
| Supramarginal gyrus | -58 | -32 | 40 | 725 | 3.3 |
| Supramarginal gyrus | 60 | -26 | 38 | 593 | 3.4 |
| Middle cingulate gyrus | 4 | 10 | 42 | 85 | 2.3 |
| Middle frontal gyrus | -36 | 52 | 18 | 541 | 2.5 |
| Middle frontal gyrus | 36 | 44 | 30 | 357 | 2.3 |
| Posterior cingulate gyrus | -4 | -52 | 28 | 153 | -2.1 |
| Angular gyrus | -48 | -64 | 38 | 587 | -2.4 |
| Angular gyrus | 54 | -62 | 38 | 60 | -2.1 |
| Orbital med. frontal gyrus | -2 | 28 | -30 | 114 | -1.9 |
| Orbital inferior frontal gyrus | -44 | 34 | -14 | 21 | -2.1 |
| Middle temporal gyrus | -60 | -42 | -2 | 235 | -2.1 |
| Inferior temporal gyrus | -62 | -10 | -22 | 208 | -2.1 |
| **CAP 7** |  |  |  |  |  |
| Postcentral gyrus | -30 | -34 | 68 | 2627 | 3.7 |
| Postcentral gyrus | 32 | -36 | 62 | 2341 | 3.4 |
| Précentral gyrus | -32 | -24 | 64 |  | 3.7 |
| Precentral gyrus | 34 | -24 | 64 |  | 3.4 |
| Supplementary motor area | -4 | 18 | 50 |  | 3.7 |
| Supplementary motor area | 10 | 4 | 42 |  | 3.4 |
| Middle cingulate gyrus | 2 | -10 | 48 | 50 | 2.1 |
| Thalamus | 4 | -22 | 2 | 58 | -2.5 |

The lmm-based ANOVA of the temporal parameters of CAPS revealed that CAP3 showed higher occurrences and higher entry rates in the “post neg. affect” resting blocks (Figure 3B), with significant differences relative to the “post neu” rest and the “post neu+task” rest blocks in direct post-hoc comparisons of the least squares means (Table 1).

**Table 1.** Results of LMMs for the analyses of occurrences (models O1 CAP3; O1 CAP4; O1 CAP7), entry rates (models E1 CAP3; E1 CAP4; E1 CAP7), and durations (models D1 CAP3; D1 CAP4; D1 CAP7), across the 5 resting conditions. Abbreviations: B= Baseline; N= post neutral; NT=post neural + task; A= post negative affect; AT= post negative affect + task. ^.^ p>0.1; * p>0.05; ** p>0.01; ***p>0.001.

| | Model | ANOVA | Estimates $\beta$ (SE) | | | | | LS means difference |
| --- | --- | --- | --- | --- | --- | --- | --- | --- |
|  |  |  | B | N | NT | A | AT |  |
| Occurrences | O1 CAP3 | F(4,72)=2.4. p=.05 | 15.6 (2.0) | 12.5 (2.0) | 11.8 (2.0) | 18.5 (2.0) | 14.0 (2.0) | A > N *<br>A > NT ** |
|  | O1 CAP4 | F(4,72)=2.5. p=.05 | 11.1 (2.0) | 14.1 (2.0) | 13.8 (2.0) | 13.8 (2.0) | 18.6 (2.0) | AT > B ***<br>AT > N *<br>AT > NT * |
|  | O1 CAP7 | F(4,72)=2.4. p=.02 | 7.5 (2.6) | 11.2 (2.6) | 13.0 (2.6) | 5.5 (2.6) | 7.7 (2.6) | NT > A **<br>N > A *<br>NT > AT *<br>NT > B * |
| Entry rate | E1 CAP3 | F(4,72)=2.8. p=.03 | 15.6 (1.3) | 11.3 (1.3) | 13.8 (1.3) | 17.3 (1.3) | 13.8 (1.3) | B > N *<br>A > NT **<br>A > N * |
|  | E1 CAP4 | F(4,72)=0.8. p=.4 | 12.1 (1.3) | 13.8 (1.3) | 15.0 (1.3) | 13.3 (1.3) | 15.4 (1.3) |  |
|  | E1 CAP7 | F(4,72)=3.8. p=.01 | 8.0 (2.2) | 12.1 (2.2) | 12.9 (2.2) | 5.6 (2.2) | 9.2 (2.2) | N > A **<br>TN > A **<br>TN > B * |
| Duration | D1 CAP3 | F(4,72)=0.8. p=.4 | 3.4 (.3) | 3.1 (.3) | 2.9 (.3) | 3.6 (.3) | 3.5 (.3) |  |
|  | D1 CAP4 | F(4,72)=2.4. p=.05 | 3.0 (.3) | 3.1 (.3) | 2.6 (.3) | 3.5 (.3) | 4.1 (.3) | AT > N *<br>AT > NT **<br>AT > A ** |
|  | D1 CAP7 | F(4,72)=1.6. p=.4 | 1.5 (.4) | 2.0 (.4) | 2.4 (.4) | 2.3 (.4) | 1.7 (.4) |  |

Similarly, a subsequent lmm analysis without the baseline condition (2 emotion contexts x 2 preceding event conditions) yielded a main effect of “emotion context”, due to overall higher occurrences and higher entry rates at rest after negative movies compared to neutral movies condition (Table 2). There was no effect of preceding event (cognitive task or movie alone).

A second resting brain network modulated by the preceding events was CAP4. This CAP implicated bilateral fronto-parietal areas bearing similarities with the dorsal attention network (Fox et al. 2006) and the dorsal anterior cingulate cortex (dACC), co-activating with medial parietal areas around the precuneus. Conversely, co-deactive areas included parts of the social cognition network (Pillemer et al. 2017; Puhan et al. 2017), such as the bilateral superior temporal sulcus (STS) and temporo-parietal junction (TPJ) the left orbital inferior frontal gyrus (IFG), as well as lateral occipital visual regions (Figure 4A, Table 3).

**Figure 4.**
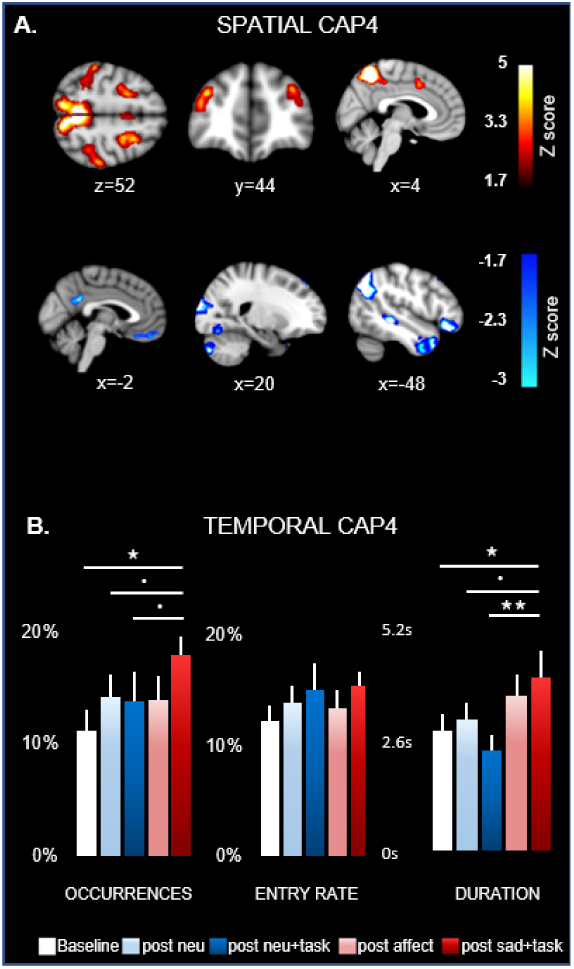
**A)** Temporal analysis of CAP4: occurrence rate (left), entry rate (middle), and duration (right) across all participants are shown for all conditions. Stars indicate significant differences between conditions (see Results section and table 3). Error bars indicate standard errors of the means (SEM). **B)** Spatial analysis of CAP4. Hot colors (above) represent areas with transient positive (co-)activation with the precuneus seed, while cold colors (below) represent areas with transient negative (de-) activation with precuneus. (see Supplementary table S7 for a further description). Slice coordinates are provided in MNI space. Brain regions were observed at Z =< −1.7 (p<.05).

The analysis of temporal metrics across all conditions revealed predominant differences in the occurrence rates and durations of this CAP during the “post neg. affect+task” resting blocks compared to both the “baseline” and the “post neu+task” resting blocks (Figure 4B, Table 1). The 2×2 analysis without baseline further showed a main effect of context (neg. affect vs. neutral) for the “duration” metric, reflecting more prolonged dwell times of CAP4 after negative affect movies (Table 2) but no other effects. There was no main effect of preceding event type (post movies vs. post cognitive task) for any of the temporal metrics.

The third network was CAP7 and comprised co-activation patterns in both lateral brain regions (postcentral and precentral gyri) and more limited medial frontal regions (anterior cingulate cortex and supplementary motor cortex). Co-deactivation patterns selectively involved the thalamus (in medial dorsal and ventral anterior regions) region (Figure 5A, Table 3). The temporal parameters showed significant differences in the occurrence and entry rates during the “post neu” and “post neu+task” conditions relative to all other conditions, i.e., “baseline”, “post neg. affect”, and “post neg. affect+task” blocks (see Figure 5B, Table 1). Consequently, significant main effects were observed in the 2×2 analysis for the same metrics (occurrences and entry rates), indicating that this specific CAP was more present after exposure to “neutral” than “negative affect” movie content (Table 2).

**Figure 5.**
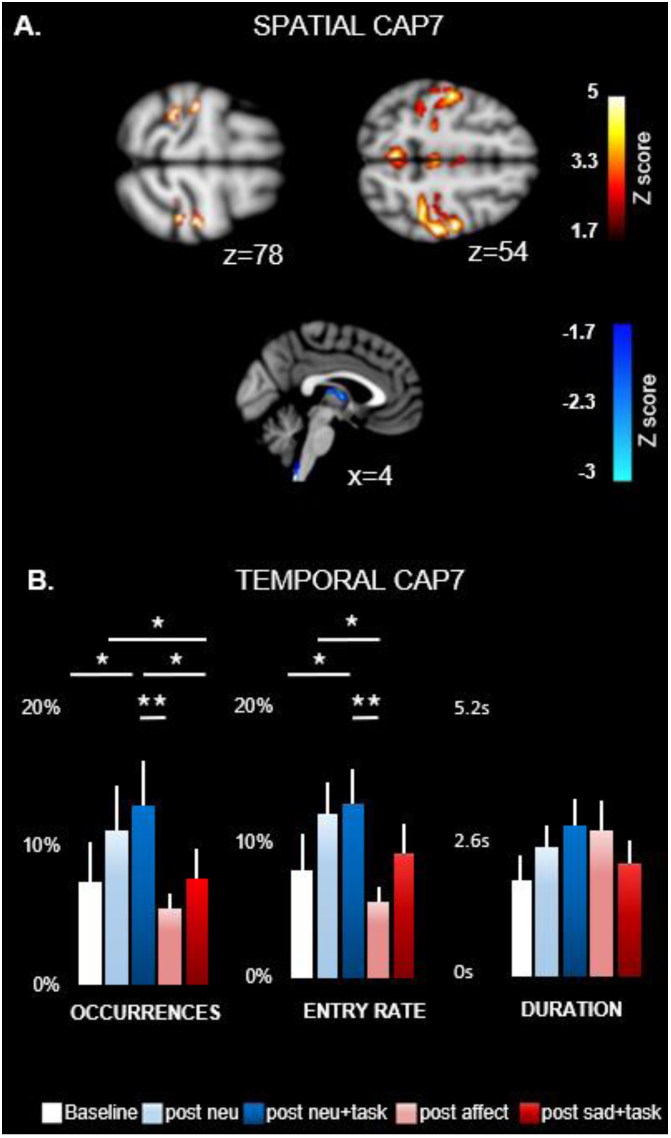
**A)** Temporal analysis of CAP7: occurrence rate (left), entry rate (middle), and duration (right) across all participants are shown for all conditions. Stars indicate significant differences between conditions (see Results section and table 3). Error bars indicate standard errors of the means (SEM). **B)** Spatial analysis of CAP4. Hot colors (above) represent areas with transient positive (co-)activation with the precuneus seed, while cold colors (below) represent areas with transient negative (de-) activation with precuneus (see Supplementary table S7 for a further description). Slice coordinates are provided in MNI space. Brain regions were observed at Z =< −1.7 (p<.05).

### Linking brain CAPs during rest to affective and behavioral measures

To better understand the functional significance of the lasting effects of emotion and cognitive control load on brain activity dynamics, we examined how the expression of the relevant CAPs above was related to behavioral measures of task performance and subjective affect following the different movie conditions. To this aim, we tested for correlations between the temporal metrics of each CAP in each participant and either the affective rating scores obtained after movies (using the PANAS), or the reaction time indices of interference measured during cognitive tasks. These analyses converged with our main results for these temporal parameters in the previous section and revealed a different impact of emotional movie content and task performance on the different CAPs.

For CAP3, individual occurrence rates during the “post neg. affect” resting condition were significantly correlated with affective scores in the negative PANAS scale from post-experiment ratings (r(18)= 0.42, p< .03). Although this correlation did not survive a correction for multiple comparisons, it was highly selective for this brain CAP and this behavioral index (Suppl. Table S5. A.), suggesting that this network configuration was not only more frequent following movies involving a negative valence (see Figure 3A), but also associated with more negative affect. Furthermore, a parametric GLM analysis (on contrast of CAP3 for “post negative” > “post neutral” conditions), using the negative PANAS (post-experiment ratings) as linear covariate, revealed higher co-activation of the ventromedial prefrontal cortex, posterior cingulate cortex, and cuneus, correlating with higher negative ratings in the sad context (Figure 6. A). Conversely, similar regression analyses with the behavioral interference scores showed no correlation with temporal metrics of CAP3, and no modulation of its spatial configuration as a function of cognitive performance.

**Figure 6.**
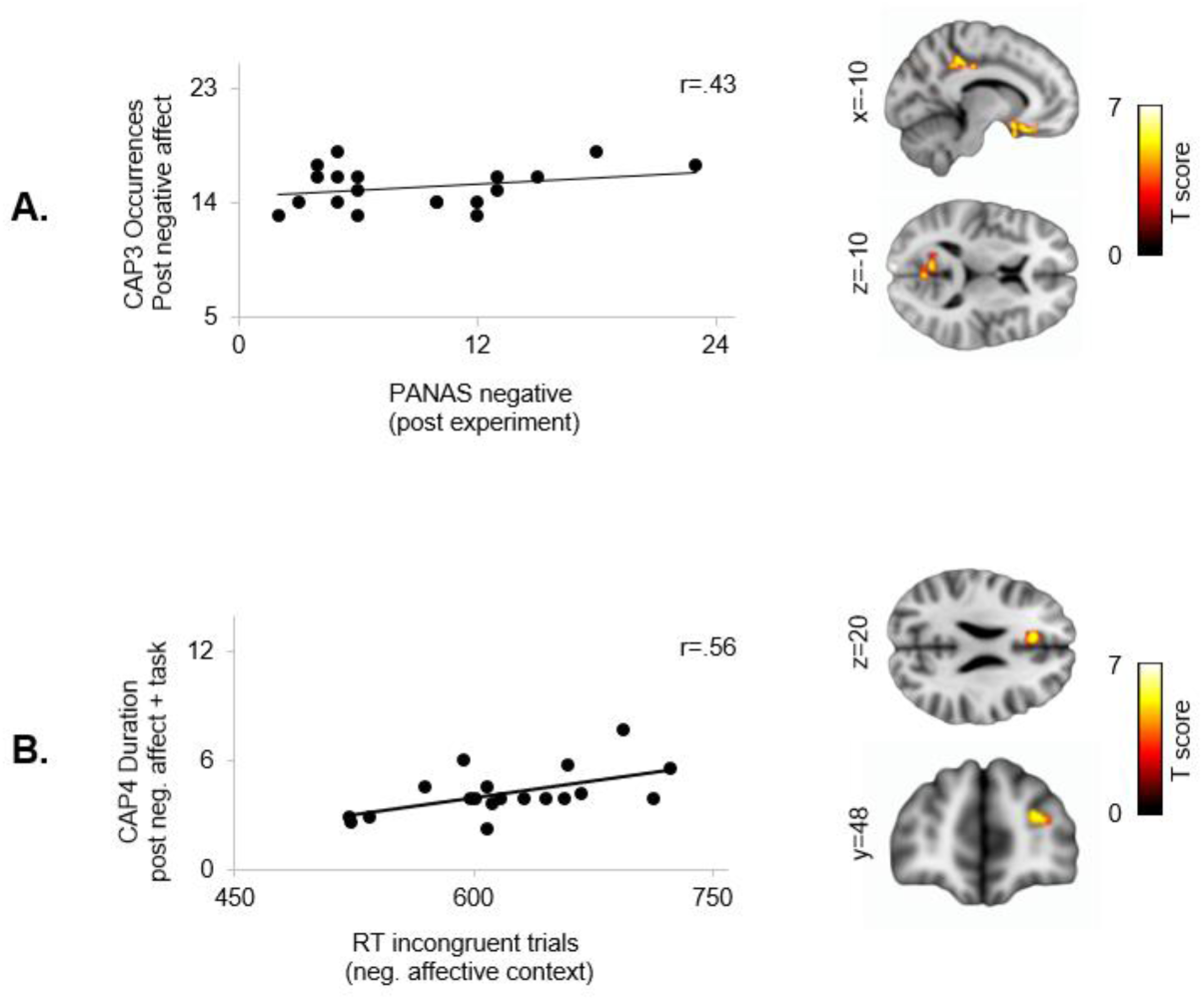
**A.** Pearson correlation coefficient scatterplots illustrating a significant linear relationship between negative PANAS scores and occurrence rate of CAP3 in the “post neg. affect” context. Right. Parametric analysis of the spatial CAP3 map (contrast “post negative affect > post neutral”) with the PANAS negative scores (post experiment) as a regressor. The ventromedial prefrontal cortex (−12 28 −18), posterior cingulate cortex (−10 38 44), and cuneus (−10 −62 10) exhibited higher co-activation within CAP3 in participants who showed more negative affective states in post-experiment ratings (p < .001 uncorrected; K=> 40). **B.** Pearson correlation coefficient scatterplots illustrating a significant linear relationship of CAP4 duration with incongruence effect in RTs during cognitive control tasks. Right. Parametric analyses of the spatial CAP4 map (contrast “post neg. affect + task” > “post neutral + task”) with incongruent RTs from the neg. affective context as a regressor. The right MFG (34 18 26) and the left ACC (−10 28 22) were significantly more co-activated within CAP4 network in participants who exhibited stronger RT slowing in this condition (p < .05 FWE corrected at cluster level; K=> 48).

Only CAP3 in the “post neg. affect” context showed a significant positive association with negative PANAS scores post-experiment. **B.** Correlations between incongruent trials (RT) in cognitive control tasks (neg. affect and neutral contexts) and duration (average) of all CAPs during the resting conditions “post neg. affect+task” and “post neutral affect+task” respectively. Only CAP4 in the “post neg. affect+task” context showed a significant positive association to the incongruent RTs in the same emotional context. (‘*’ p < 0.05, corrected for multiple comparisons).

For CAP4, the duration of the “post neg. affect+task” condition was found to strongly correlate with the incongruence costs in RTs during the cognitive control tasks, particularly in the “negative affect” context (r(18)= .56, p<.05, corrected for multiple comparisons). These data suggest that slower responding and/or greater difficulty in executive control during the task was associated with increased expression of this network in subsequent rest condition. Conversely, temporal parameters of CAP4 did not correlate with the RT costs on incongruent trials from the neutral context [r(18)=-.02, p<1] (see Suppl. Table S5. B). Remarkably, the (emotionally-modulated) RT of incongruent trials was also associated with a modulation of the spatial configuration of CAP4. Specifically, a GLM parametric analysis of CAP4 (contrast “post neg. affect+task” > “post neu+task” resting blocks), using the incongruent RT cost (from the negative context) as covariate, revealed a positive relationship with engagement of the right MFG and left ACC with increasing interference (p<.05 FWE corrected at cluster level, see Figure 6. B.). No correlations were found for CAP7 with either affective or behavioral performance scores, both for temporal parameters and spatial maps.

### Time-series variability analysis across conditions

A distribution analysis on the occurrence rates of CAPs 3, 4 and 7 over the 5 minute period of resting blocks (divided in one-minute time-bins) across the different affective contexts (see Figure S4) yielded an unequal time-course of the occurrences in the “post neg. affect” [*X*^2^ (4, N=361) = 13.8, p = .008] and “post affect+task” conditions [*X*^2^ (4, N=396) = 10.77, p = .02]. A post-hoc test of the same *X*^2^ distribution analysis revealed significant differences for CAP3, with increasing occurrence rate at minute four (in the post neg condition). CAP4 also showed a significant increase at minute three (in the “post affect sad” condition), while a modest but significant difference was observed for CAP7 at minute 1, relative to the following time bins (p < 0.05; see Suppl. Figure S4).

## Discussion

### Lingering effects of emotion and cognitive load on subsequent resting brain activity

Our findings extend previous work (Eryilmaz et al. 2011) by showing that resting brain activity is markedly altered by preceding events, particularly their affective significance. We show that exposure to negatively valenced (sad) movies did not only elicit differential neural and physiological (pupillary) activity during movie watching, but also modulated activity patterns during subsequent rest periods, in both condition-specific and network-specific manners. Negative emotions elicited by movies also influenced behavioral performance in cognitive control tasks (Etkin et al. 2011; van Steenbergen 2015), as well as the impact produced by these tasks themselves on resting brain activity. These changes were characterized by different temporal metrics of dynamic functional connectivity as defined by a data-driven analysis of co-activation patterns (CAPs), and highlighted three distinct networks (CAPs 3, 4 and 7) whose expression at rest was modulated by different affective and task conditions. Remarkably, these changes in network dynamics correlated with specific behavioral indices of subjective affect and behavioral performance, further supporting their functional relevance. Taken together, our findings demonstrate that emotional episodes may produce lasting “inertia” effects on brain networks, presumably reflecting regulatory mechanisms serving to restore normal homeostatic conditions as well as promoting adaptive behaviors in response to environmental challenges. Below we discuss the main CAPs associated with such changes and their possible functional significance.

### Affective aftermath reflected in CAP3

The occurrence and entry rates of CAP3 at rest were specifically increased in post-affective contexts (after sad relative to neutral movies), particularly when rest followed immediately the movies (rather than after the intervening cognitive task). CAP3 comprised several components of the DMN (Fig 3. Table 3). Apart from being normally active at rest, DMN areas are involved in emotion processing, self-referential mental activity and introspection, as well as autobiographical memory (Kober et al. 2008; Liemburg et al. 2012; Engen et al. 2017). DMN changes have also been reported in mood disorders (Sheline et al. 2009; Liemburg et al. 2012; Hyett et al. 2015; Lanius et al. 2015; Song et al. 2016), with DMN connectivity often regarded as a biomarker for depressive rumination (Whitfield-gabrieli and Ford 2012; Sambataro et al. 2014; Belleau et al. 2015; Hamilton et al. 2016). In line with this, we found that the occurrence rates of CAP3 in resting blocks in the post neg. affective condition (after the first movie in the affective context) and the contribution of VMPFC to this network both predicted more negative subjective affective state in post-experiment ratings. These findings suggest that the functional dynamics of CAP3 network may be critically involved in emotion regulation and self-referential processes engaged by sad information.

Simultaneously, CAP3 comprised regions from the salience network (Seeley et al. 2007) that co-deactivated with the precuneus during rest, including the anterior cingulate and insula, which have been associated to both emotional processing and self-monitoring (Lindquist et al. 2012, 2015; Touroutoglou et al. 2012; Duerden et al. 2013). Such anticorrelated DMN-SN organization (Fox et al. 2005) in the aftermath of negative emotional events is consistent with previous studies reporting that the generation of affective states may involve a coordinated recruitment of DMN and SN regions, each mediating dissociable emotion component processes (Engen et al. 2017). In this anticorrelated scheme, dynamic interactions between these networks might subserve shifts from externally oriented attention to internally oriented or self-related cognition (Menon and Uddin 2010). The co-deactive SN could thus amplify interoceptive signals and subjective emotion feelings (Chang et al. 2013) through their interaction with self-reference representations elaborated in DMN areas, implying a distinctive dynamics of connectivity of each network component with the precuneus, and ultimately contributing to conscious emotional experiences.

More generally, such DMN-SN push-pull relationship, and its enhancement in affective contexts add novel support to the notion that adaptive responses to acute stressors may imply a dynamic reconfiguration of large-scale brain networks to reallocate processing resources according to current behavioral needs, with a shift between affective and cognitive functions mediated by the SN activity as well as various neuromodulatory circuits (Hermans et al. 2014). At the functional level, this push-pull balance is also consistent with common consequences of negative emotions and stress on selective attention and thought content, as typically observed for ruminations that involve the repetitive retrieval of aversive and threat information about the self and past events that is exacerbated by negative contextual cues (Trapnell and Campbell 1999; Piguet et al. 2014). Overall, we conclude that both the temporal and spatial features of CAP3 supports a functional inertia in the dynamics of brain networks following negative (sad) events, overlapping with components of the DMN and SN, and intimately connected to subjective emotional experience as well as adaptive emotion regulation mechanisms.

### Carry-over and affective modulation of cognitive control reflected in CAP4

CAP4 showed distinctive increases in occurrence rate and duration during the “post neg. affect task” rest relative to other conditions (Figure 4). In contrast, exposure to sad movies alone did not produce reliable changes in this network. Anatomically, CAP4 comprised regions from the dorsal attention network, responsible for selective attention processing (Fox et al. 2006), including frontoparietal areas involved in executive control (Shulman et al. 2010; Ptak 2012) and dorsal ACC which is associated with the maintenance of alertness (Sadaghiani and D’Esposito 2015) as well as response conflict monitoring (Botvinick et al. 2001). This network thus partly overlaps with the so-called “cognitive control system”, a highly interconnected set of internally differentiated but reciprocally interactive units implicated in goal-directed behaviors (Cole and Schneider 2007; Vincent et al. 2008; Yeo et al. 2011). Given its modulation by the just preceding cognitive task, CAP4 might constitute a functional fingerprint of higher attentional load that extends into resting state periods after effortful task performance.

Nevertheless, this functional pattern was also amplified by the negative affective context since no similar effect was observed after task performance after neutral movies. Thus, while negative affective information itself determined changes in CAP3 during subsequent rest, attentional demands from the cognitive control tasks led to different changes in CAP4, but these effects were observed only in the “post affect +task” rest blocks, i.e., when the cognitive task was performed in a negative/sad context. Accordingly, the increased occurrence rate of CAP4 and its connectivity with medial and lateral prefrontal cortices during rest were significantly correlated with the magnitude of response interference (incongruent vs congruent) during the cognitive tasks, demonstrating a direct relationship between cognitive effort and subsequent CAP4 activity.

More broadly, these findings suggests that both affective and cognitive events can have a prolonged impact on functional brain states, and furthermore show that affective events may act not only directly on resting connectivity dynamics (as shown by modulation of CAP3 network in the “post-affect” condition) but also indirectly through a modulation of cognitive processes (as shown by modulation of CAP4 in the “post-affect+task” condition). Indeed, response conflict indices during our cognitive control task (RT cost and error rate on incongruent trials) were exacerbated after negative compared to neutral movies. These data add to previous research indicating that negative emotions are associated with decreased problem-solving abilities and impaired execution of instrumental behaviors (Nolen-Hoeksema 1991, 2015; Bonanno et al. 2008). Here we show that such effects may imply a durable re-organization and modulation of large-scale network connectivity that is apparent even during resting state and presumably impact on the mobilization of efficient neural resources during cognitive task performance. Further research is needed to more precisely compare emotion-induced changes in brain connectivity patterns during rest and during different cognitive tasks.

A co-deactive network was also observed in CAP4, with areas often implicated in social cognition such as STS, TPJ, temporal poles, PCC and vmPFC/OFC (Pillemer et al. 2017). While vmPFC/OFC is strongly involved in emotion regulation and value-based decision making (Ochsner et al. 2012; Sinha et al. 2016; Fox et al. 2018), PCC is thought to play key role in future self-related thinking and emotional contextual processing (Buckner and Carroll 2007), including moral reasoning (Greene 2001). These effects may reflect a relative disengagement of social information processing and empathic responding normally evoked after negative valenced movies but partly suppressed after performance of a difficult cognitive task. It is also noteworthy some of these areas were co-active with CAP3, when rest directly followed the negatively valenced movies. However, further analyses investigating the exact functional relationships between mental processes engaged at rest in different contexts and concomitant brain co-activation patterns will be necessary to support our interpretations.

### CAP7

A very different effect on connectivity dynamics was found for CAP7. Unlike CAP3 and CAP4, this network was functionally enhanced during rest periods in the neutral context. In addition, CAP7 comprised a thalamo-cortical network centered on sensorimotor areas (Ribary et al. 1991; Llinás et al. 1998; Ribary 2005), unrelated to higher-level cognitive or affective functions associated with other CAPs. These findings may be consistent with a role of thalamo-cortical circuits in sensory gating (McCormick and Bal 1994) as well as in regulating arousal or wakefulness (Liu, De Zwart, et al. 2018). Moreover, co-activation patterns in thalamus and postcentral cortex have been associated with cognitive load during working memory and visual attention tasks (Tomasi et al. 2006). Modulation of CAP7 may therefore reflect more externally oriented to sensory and “cold” cognitive information during rest after exposure to neutral movies (as opposed to more introspective and “warm” socio-affective processing after negatively affective movies). While this interpretation also remains speculative, the distinctive functional profile of CAP7 further highlights the selectivity of changes in connectivity dynamics induced by affective and cognitive challenges.

### Limitations and future research

A potential limitation of our study is that we included only female participants, potentially precluding generalization to males. This gender selection was based on pilot results indicating that female participants were more easily engaged with movies from the affective context, than males. Possibly because women assume more easily a first-person or empathetic perspective during the movie watching. Especially our clips featured female characters facing difficult situations that might easily occur in everyday life. Accordingly, emotional induction with social material is more reliable in female and similar selection procedure have been used in other studies (Vrtička et al. 2012). Importantly, we carefully controlled for hormonal status although using clinical interview rather than direct biological measurements. Future research would be valuable to compare the present results with those in male participants and investigate any link with individual abilities in emotion regulation, empathy, or social skills.

Another limitation is the relative short time of rest blocks (5 min). Many resting state fMRI studies employ longer durations (8-10 min), but our paradigm design required briefer sessions to avoid discomfort and boredom, which could have altered our results. Other post-emotion rest studies (Eryilmaz et al. 2011, 2014) found reliable changes with even brief rest periods (<2 min). However, seemingly inconsistent co-activation/de-activation patterns were found in the latter studies, with relative suppression of DMN activity after joyful or fearful events (Eryilmaz et al. 2011)(Harrison et al. 2008), unlike the enhanced DMN pattern of CAPs after sad movies here. On the other hand, Borchardt and colleagues (Borchardt et al. 2017) also reported increased activation of DMN-related areas in an experimental context similar to our “post neg. affect” condition. These discrepancies might be explained by differences in session duration and emotion categories, but also differences in connectivity measures, as our current approach specifically examined changes in dynamic fluctuations of co-activation patterns with the precuneus, while other studies tested for static connectivity among pre-defined areas (including DMN nodes).

Finally, these aspects might also contribute to an absence of significant differences between temporal metrics of CAP3 (post affect) and the initial baseline rest block (Fig. 2A), although comparisons of baseline with other conditions were unavoidably confounded by a time effect (since all post affect and post task blocks were necessarily obtained later). However, this time effect could not account for differences between critical subsequent conditions, all systematically counterbalanced across participants, which clearly differentiated CAP3, CAP4, and CAP7. Moreover, both CAP4 and CAP7 did exhibit significant differences relative to the baseline block, possibly driven by both condition or time factors. Lastly, while our study focused on effect induced by negative affect given previous research linking negative affect and low mood with altered cognitive functioning, it would be valuable to determine whether similar CAPs are modulated by different manipulations of emotional valence and/or cognitive challenges.

### Conclusion

Our work extends previous studies revealing a lasting impact of negative affect on brain states, beyond the transient events eliciting particular emotions, and further highlights their selective impact on functional connectivity dynamics of particular networks. In addition, we show such changes are also sensitive to the cognitive demands of preceding tasks, and negative affect further amplifies these effects at rest, beyond the direct influence of affect on task performance itself. Specifically, we find that regions linked to DMN and SN exhibit a prolonged inertia in activation patterns at rest following exposure to negative affect, predicting changes in subjective emotional status. Whereas regions linked to attentional function of FPCN and social cognition networks show antagonistic patterns determined by cognitive control performance, which correlate with task difficulty. More broadly, our study illustrates how the CAP approach provides a valuable tool to investigate temporal features of intrinsic functional brain networks. Future work probing similar CAPs and temporal dynamics in patients with mood disorders patients and affective regulation deficits might provide new insights on potential neuromarkers associated with clinical symptoms, risk, or prognosis of mental diseases.

## Notes

The authors declare no competing financial interests. This work was supported by awards from the Swiss excellence Scholarship program and the Fondation Ernst et Lucie Schmidheiny to JG; as well as by a Sinergia Grant (no. 180319) from the Swiss National Science Foundation (SNF), the Swiss Center of Affective Sciences financed by UNIGE and SNF (Grant no. 51NF40_104897) and by the Société Académique de Genève (Foremane Fund). We thank Dr. Ben Meuleman for his important suggestions regarding the statistical analysis on the CAPs data. Also, special thanks to Dr. Sevada Hovsepyan for his support during the data analysis. The analysis was carried out in the high-performance computing (HPC) BAOBAB server (https://plone.unige.ch/distic/pub/hpc/baobab_en).

## Supplementary methods

### Experimental paradigm design

Each fMRI scanning session (~62 min.) included a first 5 min resting state (baseline), followed by an emotional context period with movies containing either neutral or emotional scenes (order counterbalanced across participants). Each emotional context period included two video clips with content of the same valence (5 min per clip), two resting blocks with open eyes (5 min per block), and a cognitive task that consisted of either a Stroop or Flanker paradigm (~6 min, task type counterbalanced across emotion contexts and participants). The order of experimental blocks (rest-movie-rest-movie-task-rest) was the same in both the negative affect and neutral contexts (see Fig 1).

### Affective stimuli

Two videoclips with strong emotional content were used to modulate affective state. One movie excerpt was edited from the film “*21 Grams*” (González Iñárritu 2003), which has been used in previous mood induction studies (Hanich et al., 2014; Shiota & Levenson, 2010) and more recently in Borchardt et al. (2017). The other clip was edited from the Sophie’s Choice film (Pakula 1981), which has also been validated by previous studies in terms of valence, arousal, and type of emotions (Raz et al. 2012, 2016). These clips were selected based on several similarities regarding their content and characters, with the purpose of controlling - as best as possible - the psychological effects of naturalistic emotional stimuli. Both excerpts included young mothers as main characters, experiencing the loss of their underage children. This was intended to promote a first-person perspective in female participants, in keeping with our instruction to feel involved while watching the movies. In addition, two neutral clips of 5-minute each were taken from documentary interviews or TV pieces freely available in the internet, with verbally interacting people generally similar to sad movies but without any strong emotional aspect. The free software iMovie (https://www.apple.com/lae/imovie/) was used for editing the movies. This resulted in a set of 4 clips that were validated in a preliminary pilot study, in which a different group of healthy volunteers (n=28) rated the pleasantness, intensity, and type of emotions elicited by each movie. The final clips were compared in terms of low-level features (sound level, luminance, spatial frequency and motion), and no significant differences between both conditions were found.

### Subjective and physiological measures

Mood questionnaires assessing the participant’s affective state were given before and after the scanning session, including the state scale of the State Anxiety Inventory, STAI (Spielberger et al., 1983; Gauthier et al., 1993), and the Positive and Negative Affect Scales, PANAS (Watson et al., 1988). Additionally, participants rated their affective responses for each movie with a 9-point Likert scale combined with the self-assessment Manikin, SAM (Bradley and Lang 1994), as used in previous work (Borchardt et al., 2017; Hanich et al., 2014). These ratings included valence (“How happy/pleased or unhappy/dissatisfied are you?”, 1 = very unhappy, 9 = very happy), arousal (“How awake/aroused or calm/drowsy do you feel?”, 1 = very calm, 9 = very aroused), and subjective hedonic experience (“How pleasant or unpleasant was your own experience of the scene?”,1 = very unpleasant, 9 = very pleasant. see results in supplementary results section and Supplementary Figure S1 C.).

### Pupil recording

Pupil diameter and eye-gaze position were recorded during the entire scanning session, using an MR-compatible eye-tracking system at a sampling rate of 60Hz (Eye Track 6; Applied Science Laboratories, USA). Pupillometric data were analyzed offline after standard preprocessing. Blink artifacts were identified and removed by linear interpolation with the nearest neighboring valid data points. Trials with >25% of blink artifacts were not considered for analysis. To estimate whether emotional movies increased arousal during the movies, we averaged data points over the whole movie duration and analyzed these values using a repeated-measures ANOVA with the factor emotional context as factor (see results in Supplementary Figure S1 B.).

### Cognitive control task

To test for the effects of negative emotion on cognitive task performance, we used two classic interference paradigms (face Stroop task and number Flanker task), adjusted for fMRI compatibility. Each task comprised 80 trials (40 congruent and 40 incongruent). Individual stimulus presentation lasted 1s, followed by an inter-trial interval with a central fixation cross randomly jittered from 2 to 4.9s. Both tasks were counterbalanced across affective context conditions and across participants. Each task combined congruent (C) and incongruent (I) trials in which a central target (name in Stroop, number in Flanker) was presented for a binary classification response (male/female for Stroop, odd/even for Flanker), together with a distractor corresponding to either the same (C) or opposite (I) response. In addition, a given trial could be preceded by either the same or opposite condition (C or I trials), in a semi-random but balanced order. This resulted in 4 trial types, allowing subsequent behavioral analysis according to both current compatibility (indicated by upper-case letters C and I) and compatibility of the preceding trial (indicated by lower-case letters c and i). sec (see Fig. 1B for a schematic description)

### Face-name Stroop task

All face images in the Stroop task were obtained from the Karolinska Directed Emotional Faces, KDEF (Calvo and Lundqvist 2008). Participants were exposed to pictures of faces superimposed by names. The name and face corresponded to the same gender in 50 % of the trials (congruent stimuli). We used images of 35 male and 35 female faces (portrait format, size 6. cm x 8.5 cm). To obtain some homogeneity of visual features, all images were standardized in terms of color background (grey), color portray (black &white), and face proportion in mage (close-up). We used 70 different names from the French language and whose gender association was verified tested during preliminary piloting. All stimuli appeared in the center of a screen outside the scanner bore, reflected on a mirror placed on the head coil. Participants gave responses with an MRI-compatible device with two key buttons at the right side. The instruction was to categorize the <u>name</u> as male or female. The stimuli size and screen-participant distance were reduced in a 1:2 ratio (approx.) for the trials outside the scanner.

### Number Flanker task

The stimuli were digits 1, 2, 3, 4, 6, 7, 8, and 9, presented on gray background. They were presented with the same font size and color as names in the Stroop task. To control for negative priming, a digit could be repeated as a flanker not less than nine trials after being presented as target. The target digit appeared in the center of the screen, reflected on the head-coil mirror. Two flanker digits were presented to the left and right of the target, at a distance of 10 mm from the target. The left and right flanker digits were always the same. The subjects’ task was to categorize the central digit as odd or even, and to respond by pressing a left or right response key from the MRI-compatible device. The stimuli size and screen-participant distance were reduced in a 1:2 ratio (approx.) for the trials’ tasks given inside vs outside the scanner.

### FMRI data acquisition

Neuroimaging data were collected using a 3T Magnetom TIM Trio scanner (Siemens, Germany) and a 32 channels head-coil. The Blood Oxygenation Level Dependent (BOLD) contrast was measured using a T2*-weighted echo-planar sequence (EPI). Five hundred functional volumes of 36 axial slices each (TR/TE/flip angle = 1300ms/30ms/80°, FOV=192 mm, resolution=64×64, isotropic voxels of 3.2 mm^3^, distance factor 20%) were acquired in one single continuous scanning run. We collected a high-resolution T1-weighted anatomical image (TR/TI/TE/flip angle=1900ms/900ms/2.27ms/9°, FOV=230mm, resolution=256×256, slice thickness=0.9mm, 192 sagittal slices) at the end of the first session.

#### Preprocessing for GLM-based fMRI analysis on videoclips and tasks

Standard image preprocessing procedures were applied using SPM12 (www.fil.ion.ucl.ac.uk/spm/). Functional images from the video clips and the tasks were realigned, slice-time corrected, normalized and co-registered to individual skull stripped anatomical images comprised by the probabilistic maps of CSF (cerebrospinal fluid), gray and white matter extracted by the DARTEL Algorithm (Ashburner 2007). Data was spatially smoothed with an 8 mm Gaussian kernel.

#### GLM-based fMRI analysis

Preprocessed fMRI data from the video clips, cognitive tasks, and resting blocks were separately analyzed using the General Linear Model as implemented in SPM12. For the movie analysis, four covariates of interest were used in the regression analysis to model each clip. Regressors in these models were convolved with a standard hemodynamic response function (HRF) according to a blocked design, which was then submitted to a univariate regression analysis. Realignment parameters were also added to the design matrices of both models, to account for any residual movement confounds. In all cases, the design matrix included low-frequency drifts (cutoff frequency at 1/128 Hz). Flexible factorial analyses of variance (ANOVAs) were performed on the main contrasts of interest. All statistical analyses were carried out at the whole-brain level, with a threshold at P < 0.05, FWE corrected at whole brain level (unless specified otherwise).

#### Preprocessing for co-activation pattern (CAP) analysis of resting periods

Image processing of rest blocks included a similar procedure as in the GLM analysis. However, the kernel smoothing was reduced to 5 mm. Time series from ROIs in the white matter and cerebrospinal fluid, as well as the six affine motion parameters from realignment, were used as nuisance variables to be regressed out from the data. Furthermore, the data were band-pass filtered between 0.01 and 0.10 Hz. All image volumes with frame-wise displacement above 0.5 mm were discarded, as well as subjects with more than 50% of scrubbed frames (Power et al. 2014). A mask based on gray-matter segmentation and a probabilistic anatomical midbrain atlas were created for further analysis.

#### Parameter specification for CAP analyses

We specified the following parameters for the CAP computation. First, we established an activation threshold (T=0.7 mm) for the minimal absolute value of signal in the seed ROI (precuneus) in order to use the corresponding fMRI frames (i.e. time points) in the analysis. A second motion threshold (T_mot_=0.5mm) was also set to scrub and discard frames with excessive displacement (Power et al., 2012; Power et al., 2015). Then, we defined the percentage of positive and negative voxels in each time frame to retain for clustering at 50%, in order to improve the signal-to-noise ratio (SNR) in single fMRI volumes (Liu et al., 2013). Finally, we set at 50 the number of iterations over which our clustering was run, in order to avoid convergence to a local minimum (Dinstein et al., 1988). Consecutively, each of the CAPs obtained by clustering were transformed into spatial Z-maps by normalizing them to the standard error. This provided a quantitative measure for the degree of significance to which a CAP map values deviates from zero. Different parameters were then computed to evaluate the dynamics of each CAP.

#### Definition of temporal metrics in CAPs

First, we defined a global “occurrence rate” for each CAP as the fraction of the retained frames assigned to each cluster, computed session-wise. Second, we defined the “entry rate” of each CAP as the number of times that the seed (precuneus) entered a particular state of co-activation or co-deactivation with this CAP, computed as the sum of all the first occurrences of several successive retained time-frames for the same cluster, plus single occurrences, normalized by the total number of entries in one of the co-activation or co-deactivation for a given session. Third, we defined the “duration” of each entry rate, expressed as seconds (number of frames multiplied by the TR). Finally, spatial CAPs were computed and represented by individual activation maps, for each participant and resting condition.

## Supplementary Results

### Physiological and psychological indices of affective stimulation

The pupillary recordings showed increased dilation during negative movies (red) in comparison to neutral (blue) [F (1, 64) = 3.99, p < .01, data averaged between movies with the same emotional valence]. No differences in pupil size were found between the two neutral movies (p = 0.74) or the two affective movies (p = 0.84).

Subjective affective ratings of the negative affect and neutral movies also yielded significant differences between the two emotional contexts, relative to those made in the neutral contexts. Valence was more negative for affective than neutral clips [F (1, 72) = 3.97, p < .01] (supplementary Figure S1 B), and there was no difference between movies within the same emotional conditions (neutral movies, p = .66; affective movies, p = .84). Arousal was also higher for the affective than neutral movies [F (1, 72) = 3.96, p < .01]; no differences within each emotional condition (neutral, p = 1; negative affective, p = 0.8).

Subjective emotional experience (Likert scale from 1 to 9) was also rated as more negative after watching films with negative affect than after the neutral ones [F (1, 72) = 3.97, p < .01]. Likewise, affective questionnaires indicated significant differences in emotional state before and after the experiment. Two-tailed paired t-test comparing scores before (Pre-experimental) and after the scanning session (Post-experimental) on the PANAS showed more negative scores after the experiment [**t (**18**) =** 1.7**, p <** 0.01], and consequently, less positive scores as well [**t (**18**) >** 2.10**, p <** 0.05]. A similar pattern was observed in the STAI state scale, with higher anxiety scores after, compared to before the experiment [**t (**18**) =** 2.10**, p <** 0.001]. (See Supplementary Figure S1. E).

### GLM-based fMRI analysis

Comparing brain activation during movies in both contexts showed differential in the “neutral > neg. affective” contrast and the inverse “neg. affective > neutral”, described in Suppl. Table S1. The seed ROI selected in precuneus for the subsequent CAP analysis (as this region constitute a main hub active during resting state (Cavanna and Trimble 2006; Margulies et al. 2009) also showed increased activity in the contrast “neg. affect clips > neutral clips” (bilateral cluster with peak at 4 −52 54; p < .05, FWE-corrected for the whole level; k = 486).

### Behavioral data from cognitive tasks

A standard interference effect (I > C) was found in response time (RT) and error rate (ER) during both tasks: Flanker RT: t (18) =-8.70, P < 0.001; Flanker ER: t (18) =-3.9, P < 0.001; Stroop RT: t (18) = −9.32, P < 0.001; Stroop ER: t (36) =-2.7, P < 0.001 (see Suppl. Figure S2 A). The interference effect (I minus C) showed no significant differences between the two tasks [F (1,18) = 1.3, p = .5]; see Suppl-Figure S2 B. The interference effect was reliably found in both RT and ER following both negatively-valenced emotional clips [RT: t (18) =3.24, P < 0.001. ER: t (18) =-3.08, P < 0.01] and neutral clips [RT: t (18) = 2.12, P < 0.05. ER: t (18) =-2.28, P < 0.05], but the effect was larger in the former condition [RT: t (18) = 2.60, P < 0.01; ER: t (18) = −.34, P < 1] (see Suppl. Figure S1 D). These behavioral data confirm that incongruent trials were more demanding than congruent trials regardless of the kind of task, but this cognitive costs was amplified after exposure to sad compared to neutral clips.

#### BOLD correlates of the task performance

To examine the neural substrates recruited by cognitive control tasks, we compared fMRI activity during the Stroop and Flanker tasks (combined) using a standard GLM analysis. Consistent with previous studies (Botvinick et al. 2001; Kerns 2004; Egner and Hirsch 2005; Egner 2007; Egner et al. 2008; Rey et al. 2014), we found robust congruency effects (CE) reflecting interference (I>C trials) in several brain areas involved in conflict monitoring and selective attention control (see Supplementary table S2).

Further, differences between affective contexts were determined by testing the interaction contrast “I>C neg. affect vs. I>C neutral” (and vice versa). This revealed significant increases in right prefrontal cortex (MFG) in neg. affective condition but reduced activity in medial brain areas typically associated to the DMN. Finally, a main effect of emotion context (all trials types after neg. affect movies > all trials after neutral movies) showed increased activity in the medial and posterior bilateral insula, a region typically related to affective processing and salience monitoring (Supplementary Table 3).

---

**Supplementary Figure S1.**
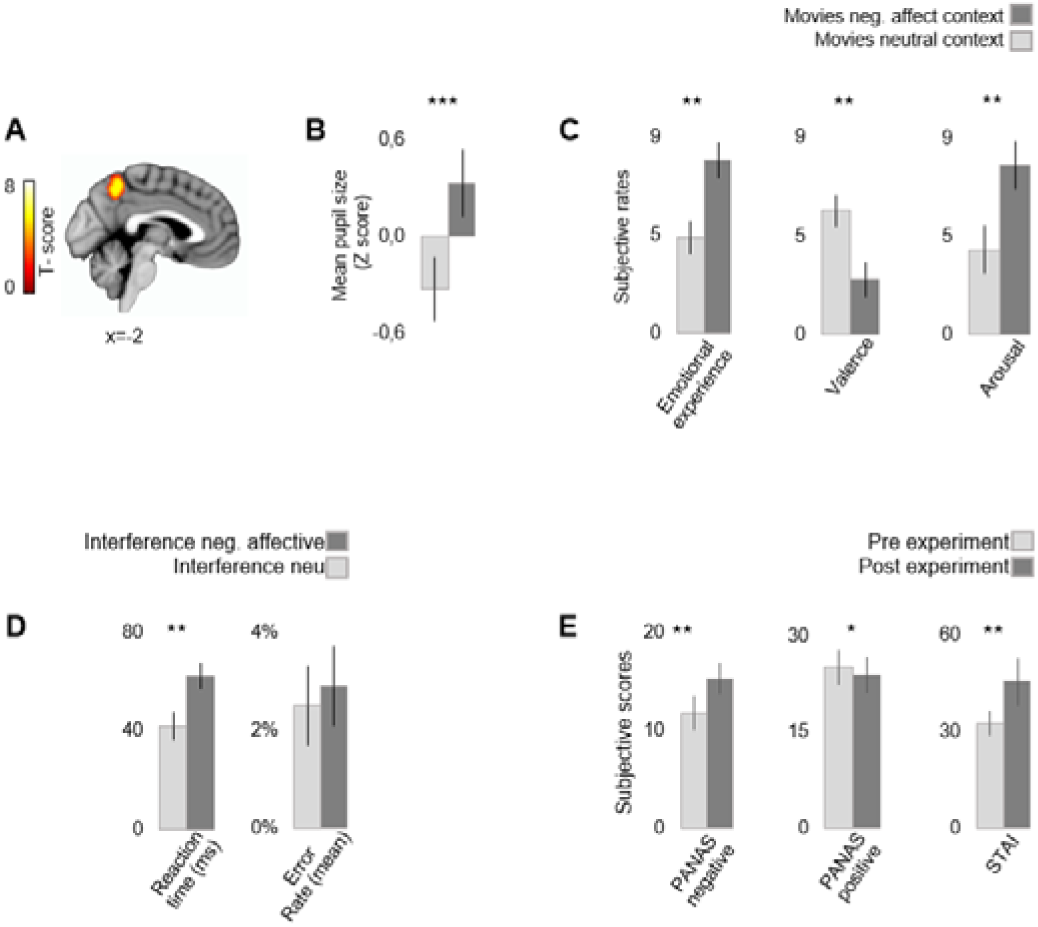
A) GLM-based Seed region in precuneus activated used for CAP analysis of resting state conditions and showing increased activity during neg. affect vs neutral movies. B) Pupillary recordings illustrating increased dilation during neg. affect compared to neutral movies. C) Subjective affective ratings of movies obtained post-scanning. D) Behavioral data from cognitive control tasks showing interference indices (performance on incongruent minus congruent trials) for each affective contexts, pooled across both cognitive tasks (for separate data on Stroop and Flanker tasks, see Fig S2). The bar plots denote significant differences only for RT, not error rate. E) Mean scores on PANAS and STAI measures for rating before a scanning session (Pre-experimental score) and after a scanning session (Post-experimental score), for both affective contexts. Whiskers stand for standard errors of means. * p>0.05; ** p>0.01; ***p>0.001.

**Supplementary Figure S2.**
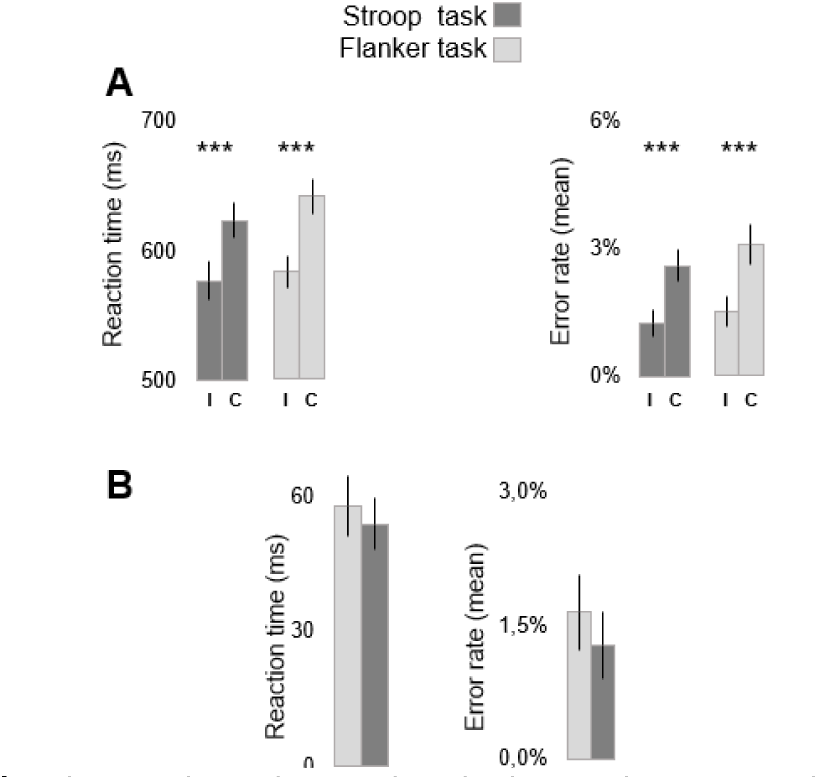
A) Behavioral results in the Flanker and Stroop tasks separately. “I” indicates the incongruent trials, and “C” congruent trials. Similar patterns are observed in both tasks. B) Magnitude of interference effect (I-C) for RTs and error rates, showing no significant differences between the two tasks. ** p>0.01; ***p>0.001.

**Supplementary Figure S3.**
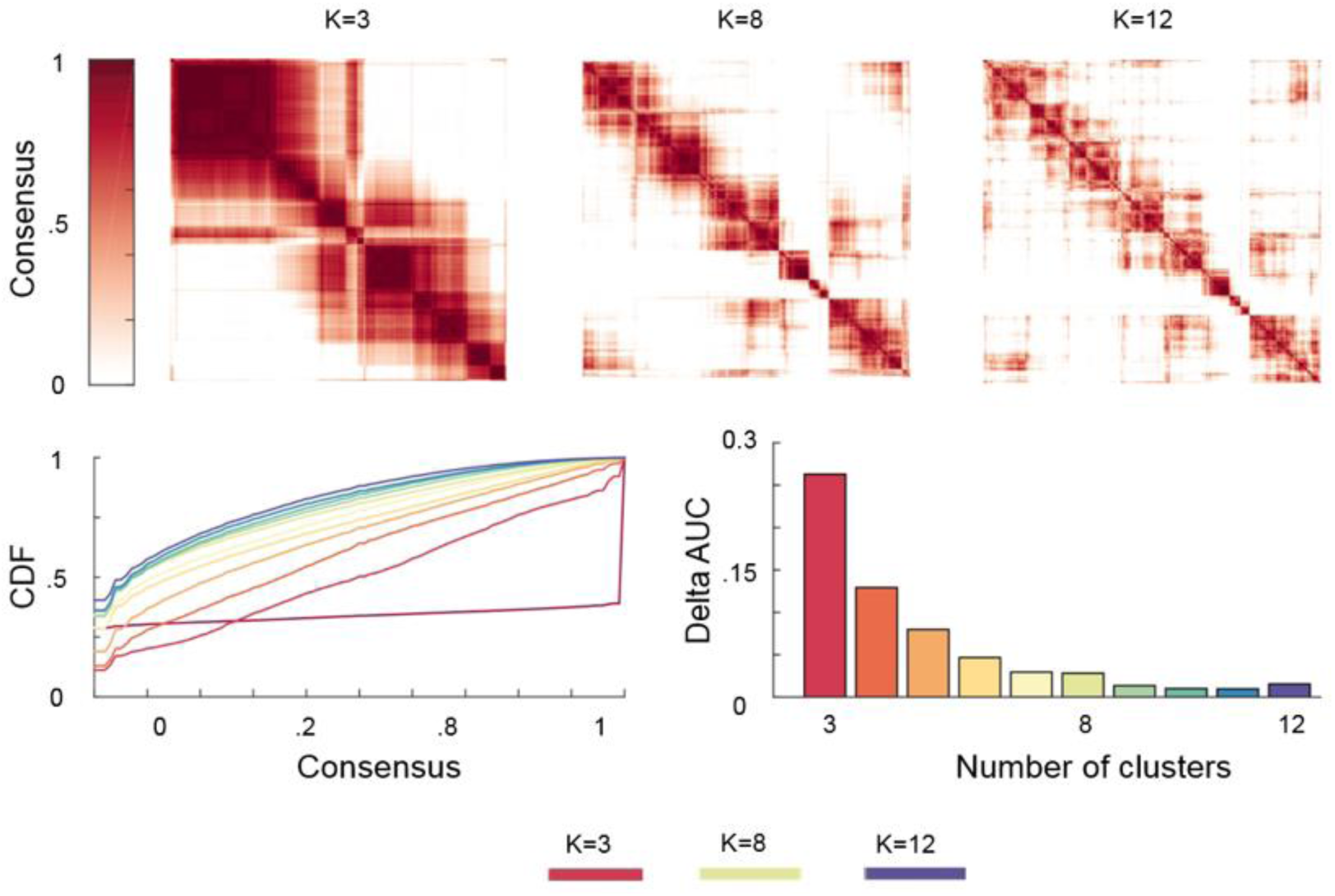
Consensus clustering analysis of CAPs identified during resting state fMRI. A consensus resampling-based clustering algorithm was implemented to obtain the number and membership (consensus) of reliable CAPs within our dataset. Parameters used to calculate a “consensus rate” between all pairs of samples included 80% of item resampling, a maximum k of 12, 50 resamplings, with Euclidean distance index and Pearson correlations as the distance measurement. Up: Heatmaps of consensus matrices for k=3, k=8, k=12, where values range from 0 (samples are never clustered together across consensus folds) to 1 (always clustered together), marked by white to dark red colors. Bottom left: Consensus Cumulative Distribution Function (CDF) of the consensus matrices from k = 2 to k = 12 (k=3, k=8, and k=12; indicated by red, laguna yellow, and purple, respectively), describing how consensus entries distribute for each case. Bottom right: Delta area under the curve plot indicating the relative change in area under the CDF curve (a larger area change implies a larger increase in the quality of clustering at the assessed k). These results provide qualitative (top matrices) and quantitative (bottom right plot) information suggesting that k = 8 is the optimal number of clusters for classifying our dataset.

**Supplementary Figure S4.**
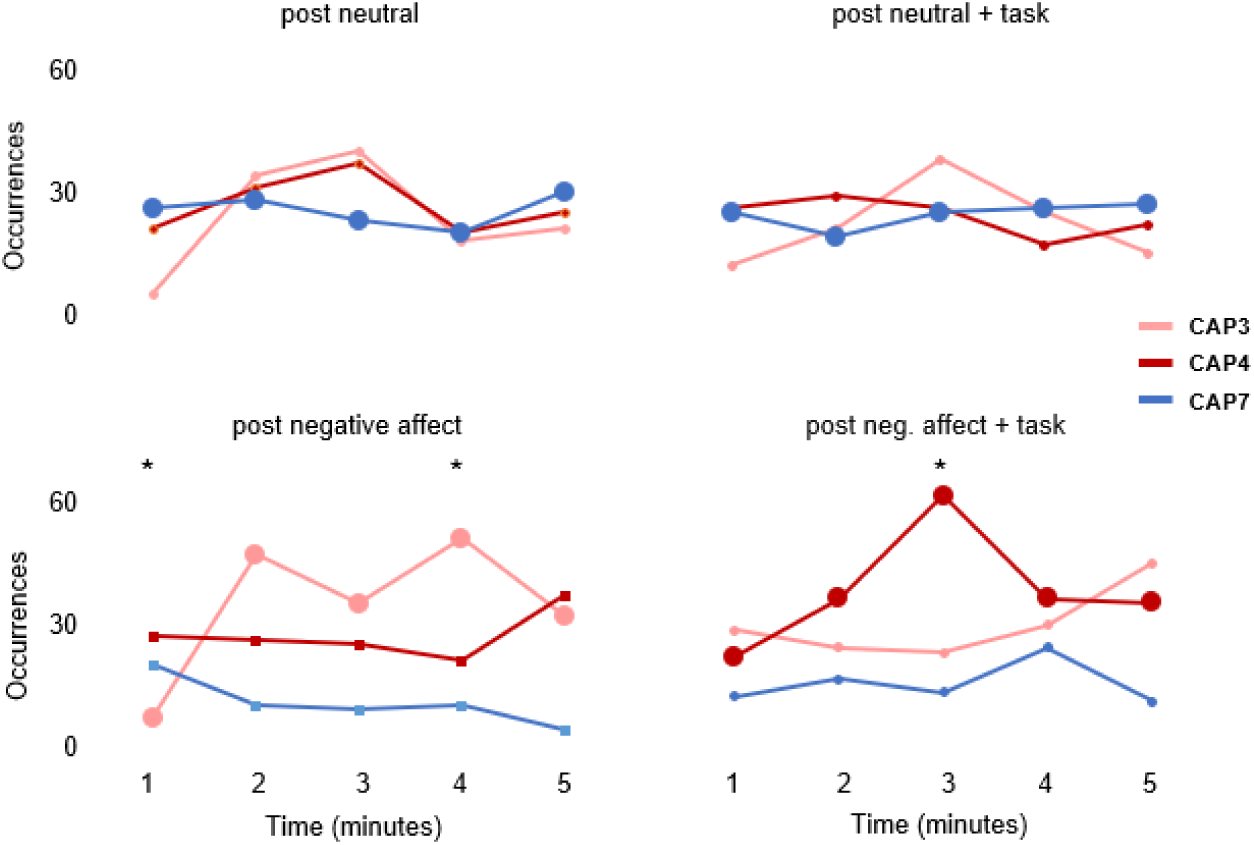
Time series analysis of occurrence rates for CAPs 3, 4, and 7 in one-minute time bins across the 4 main resting conditions as a function of preceding affective context (neg. affective vs neutral movie) and preceding events (movie with or without cognitive task). * p>0.05.

**Supplementary Figure S5.**
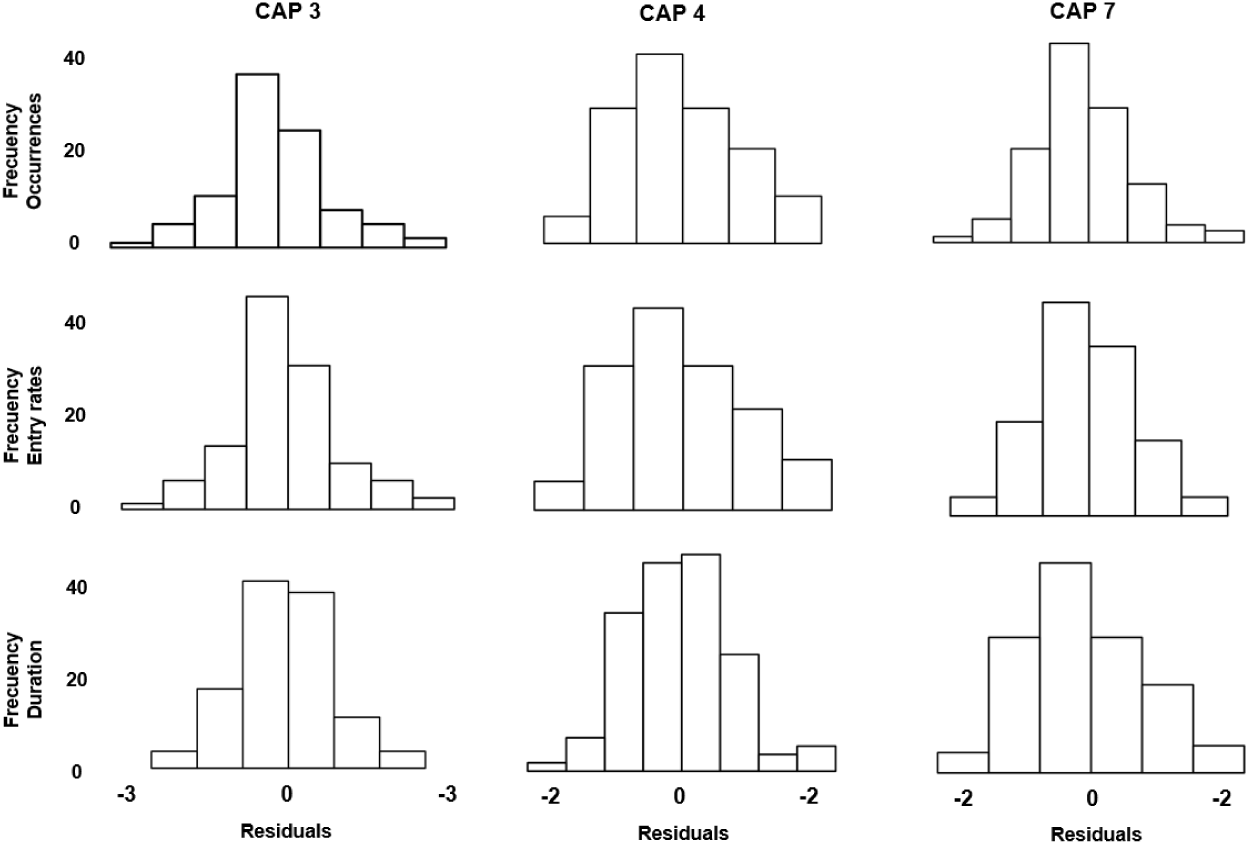
Normal probability plot of residuals. The histograms from nine different LMM analyses are relatively bell-shaped. They suggest a normal distribution of the data, and therefore no violation of the normality assumption.

**Supplementary Table S1.** Brain activation in negatively-valenced emotional movie clips vs. neutral clips. Clusters were displayed using a threshold at p < 0.05, FWE- corrected for the whole brain.

| Brain Region | Coordinates (MNI) |  |  | K | Z-score |
| --- | --- | --- | --- | --- | --- |
|  | x | y | z |  |  |
| Neutral > Negative affect |  |  |  |  |  |
| Supramarginal gyrus | -22 | -38 | -4 | 119 | 5.91 |
| Negative affect > Neutral |  |  |  |  |  |
| Inferior occipital gyrus | 52 | -70 | 2 | 839 | 9.66 |
| Inferior occipital gyrus | -46 | -78 | 6 | 784 | 7.75 |
| Precuneus | 4 | -52 | 54 | 486 | 6.50 |
| Lingual gyrus | 10 | -78 | -6 | 309 | 6.20 |

**Supplementary Table S2.** Two Mixed models (LMMs) on reaction time and error rates from the tasks. Eight regressor were included in each model, representing the kind of trial (cC=previous congruent, current congruent; iC=previous incongruent, current congruent; iI= previous incongruent, current incongruent; cI= previous congruent, current incongruent), and the context (neutral, negative affect) as fixed effects. Scores from each participant were included as random effects.

| ANOVA | | | | Estimates $\beta$ (SE) | | | | |
| --- | --- | --- | --- | --- | --- | --- | --- | --- |
|  | Current trial | Previous<br>x Current | Emotion<br>x Current | Context | cC | iC | il | cl |
| Reaction time | F(1,126)=123.7<br>p<.001 | F(1,126)=19.9<br>p< .001 | F(1,126)=4.9<br>p< .05 | Neutral | 567.6<br>(14.5) | 598.8<br>(14.5) | 619.2<br>(14.5) | 629.9<br>(14.5) |
|  |  |  |  | Negative affect | 571.7<br>(14.5) | 587.5<br>(14.5) | 619.2<br>(14.5) | 654.1<br>(14.5) |
| Error rates | F(1,126)=14.4<br>p<.001 | F(1,126)=0.2<br>p< .1 | F(1,126)=.6<br>p< .1 | Neutral | .36 (.2) | .73 (.2) | 1.1 (.2) | .95 (.2) |
|  |  |  |  | Negative affect | .42 (.2) | .68 (.2) | 1.1 (.2) | 1.1 (.2) |

**Supplementary Table S2.** Brain activation after emotionally-influenced cognitive control tasks. Up. Regions responding to the contrast “Interference Neutral > Interference Neg. affect” (p<.005, uncorrected). Center. Contrast Interference Neg. affect > Interference Neutral (p<.005, FWE corrected at cluster level). Main effect of the negative affective context > neutral context (p<.001, FWE corrected at cluster level).

| Coordinates (MNI) |  |  |  |  |  |
| --- | --- | --- | --- | --- | --- |
| Brain Region | x | y | z | K | Z-score |
| Congruent > Incongruent |  |  |  |  |  |
| Hippocampes | -22 | -38 | -4 | 119 | 5.91 |
| Incongruent > Congruent |  |  |  |  |  |
| Superior Parietal lobe | -22 | -62 | 46 | 784 | 8.04 |
| Superior Parietal lobe | 28 | -68 | 30 | 675 | 5.39 |
| Middle frontal Gyrus | -24 | 4 | 48 | 188 | 6.33 |
| Supplementary Motor C. | -80 | 20 | 48 | 250 | 5.43 |
| Inferior temporal Gyrus | -48 | -66 | -16 | 372 | 5.07 |
| Inferior temporal Gyrus | 46 | -62 | -14 | 129 | 5.01 |
| Middle Occipital Gyrus | 36 | -82 | 12 | 287 | 5.29 |
| Middle frontal gyrus | -40 | 26 | 22 | 140 | 5.81 |
| Middle frontal gyrus | 42 | 10 | 34 | 126 | 5.08 |
| Putamen | -22 | 0 | 4 | 118 | 4.73 |

**Supplementary. Table S3.**
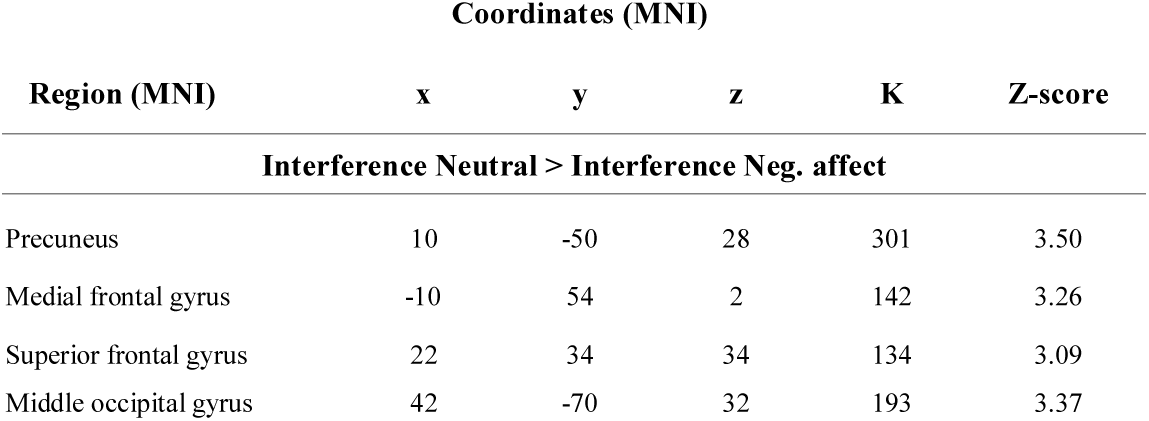
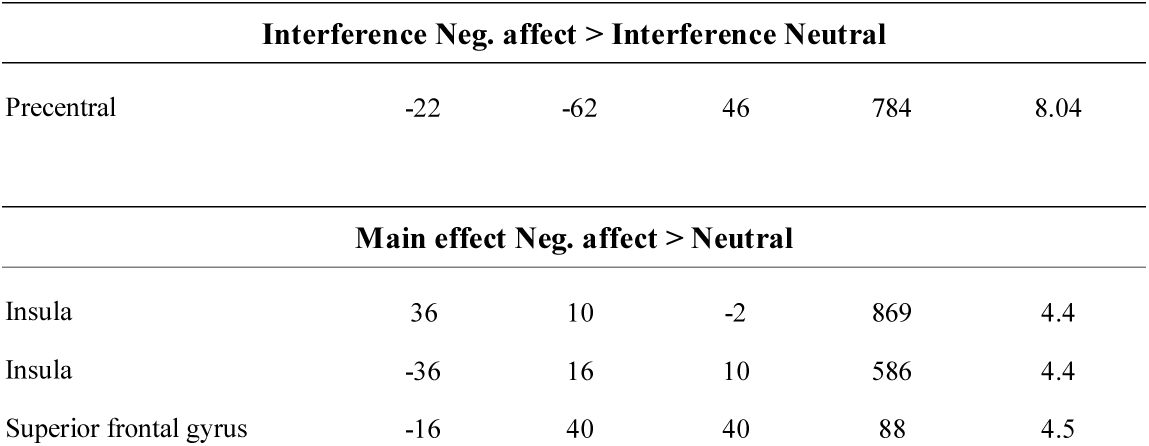
Effects of the interference effect on BOLD activity. Up. brain regions with increased activity in the contrast congruent > incongruent. Bottom. brain regions with increased activity in the contrast incongruent > congruent trials. (p < .0001, FWE-corrected at cluster level).

| Coordinates (MNI) |  |  |  |  |  |
| --- | --- | --- | --- | --- | --- |
| Region (MNI) | x | y | z | K | Z-score |
| Interference Neutral > Interference Neg. affect |  |  |  |  |  |
| Precuneus | 10 | -50 | 28 | 301 | 3.50 |
| Medial frontal gyrus | -10 | 54 | 2 | 142 | 3.26 |
| Superior frontal gyrus | 22 | 34 | 34 | 134 | 3.09 |
| Middle occipital gyrus | 42 | -70 | 32 | 193 | 3.37 |

| Interference Neg. affect > Interference Neutral |  |  |  |  |  |
| --- | --- | --- | --- | --- | --- |
| Precentral | -22 | -62 | 46 | 784 | 8.04 |

| Main effect Neg. affect > Neutral |  |  |  |  |  |
| --- | --- | --- | --- | --- | --- |
| Insula | 36 | 10 | -2 | 869 | 4.4 |
| Insula | -36 | 16 | 10 | 586 | 4.4 |
| Superior frontal gyrus | -16 | 40 | 40 | 88 | 4.5 |

**Supplementary Table S5.** **A.** Correlations between PANAS negative scores (post-experiment) and occurrence rates of all CAPs during the resting conditions “post negative affect” and “post neutral”.

|  |  |  |  |  |  |  |
| --- | --- | --- | --- | --- | --- | --- |
| **A.** | CAPs (Occurrences)<br>Post negative affect | PANAS negative<br>(post experiment) |  | CAPs (Occurrences)<br>Post neutral | PANAS negative<br>(post experiment) |  |
|  |  | **r** | **p** |  | **r** | **p** |
|  | 1 | .06 | .4 | 1 | .04 | .9 |
|  | 2 | -.06 | .5 | 2 | -.09 | .7 |
|  | 3 | **.43** | **.03** | 3 | -.36 | .1 |
|  | 4 | .09 | .3 | 4 | -.16 | .5 |
|  | 5 | .00 | .5 | 5 | .13 | .6 |
|  | 6 | -.03 | .5 | 6 | .03 | .9 |
|  | 7 | -.31 | .9 | 7 | .28 | .2 |
|  | 8 | .03 | .4 | 8 | -.15 | .6 |
| **B.** | CAPs (Duration)<br>Post neg. affect + task | RT incongruent trials<br>(neg. affective context) |  | CAPs (Duration)<br>Post neutral affect + task | RT incongruent trials<br>(neg. affective context) |  |
|  |  | **r** | **p** |  | **r** | **p** |
|  | 1 | -.37 | .9 | 1 | .31 | .9 |
|  | 2 | -.03 | .5 | 2 | -.50 | .01 |
|  | 3 | -.16 | .7 | 3 | -.11 | .6 |
|  | 4 | **.56** | **.04\*** | 4 | -.02 | .9 |
|  | 5 | .00 | .3 | 5 | .18 | .46 |
|  | 6 | .12 | .3 | 6 | .17 | .4 |
|  | 7 | -.37 | .9 | 7 | -.03 | .9 |
|  | 8 | -.05 | .5 | 8 | .08 | .7 |

